# The integration of prosody and semantics in non-literal speech: A voxel-wise encoding model approach using large language models

**DOI:** 10.64898/2026.08.21.746185

**Authors:** Wittmann Adrien, Ceravolo Leonardo, Grandjean Didier

**Affiliations:** Neuroscience of Emotions and Affective Dynamics lab, Swiss Center for Affective Sciences, Department of Psychology and Educational Sciences, University of Geneva, Geneva, Switzerland

## Abstract

Irony and sarcasm are complex forms of non-literal language that hinge on a misalignment between surface meaning and speaker intent, requiring listeners to integrate contextual, semantic, and prosodic cues. While prior neuroimaging studies have implicated a broad network—including the temporal cortex, the inferior frontal gyrus, and the medial prefrontal cortex—in the comprehension of ironic and sarcastic speech, the precise neural mechanisms underlying the integration of semantic and prosodic information remain unclear. In the present study, we addressed this gap by employing voxel-wise encoding models to systematically identify brain regions specifically involved in combining prosodic and semantic cues during non-literal language comprehension. Participants listened to naturalistic auditory dialogues in which both discourse context and target utterance semantics and prosody were systematically manipulated. We derived custom text embeddings using transformer-based models to capture context-sensitive semantic representations of ironic statements, alongside acoustic features characterizing affective prosody. Ridge regression models were fitted to predict BOLD responses at the voxel level using semantic, prosodic, and combined features, and we identified integration as voxels in which each modality contributed predictive information beyond the other, using a permutation-based conjunction test. The regions integrating prosody and semantics depended on whether discourse context was modeled: integration was confined to the bilateral temporal speech cortex when statements were encoded in isolation, but additionally engaged the left inferior frontal gyrus pars orbitalis (IFGorb) when each statement was weighted by its relevance to the preceding context. These findings indicate that the left IFGorb integrates prosody with context-dependent meaning, engaging beyond the temporal speech cortex specifically when comprehension requires combining semantic, prosodic, and contextual cues—as in irony and sarcasm.

**Author summary:** In our daily conversations, people often say the opposite of what their words literally mean. When someone is being ironic or sarcastic, listeners rely not only on what is said but also on how it is said—the tone of voice—and on the broader context. We wanted to understand how the brain brings these pieces together to recover a speaker’s true meaning. Using artificial intelligence tools, we modeled the meaning of each sentence and gave more weight to the words most strongly linked to the earlier context. We combined these representations of meaning with measurements of vocal tone and tested how well they predicted brain activity while people listened to short dialogues. By comparing models that used only meaning, only tone, or both, we pinpointed regions that respond specifically to the combination of the two. When each sentence was modeled on its own, only speech regions in the temporal lobe combined tone and meaning. But once we let the context reshape a sentence’s meaning, a higher-level region in the left frontal lobe also came into play. This suggests that interpreting irony and sarcasm relies on a frontal region that blends tone with meaning after context has shaped it.

## Introduction

Irony and sarcasm are types of non-literal language in which the speaker’s intended meaning diverges sharply from the literal content of their words [1–3]. At their core, these communicative forms rely on a clash between surface meaning and contextual intent. Irony often functions as a rhetorical strategy to highlight incongruity, sometimes with a humorous or reflective aim. In particular, ironic praise involves using seemingly critical language to convey admiration or affection. For example, saying “I see that you are again at the bottom of the class” to a friend who just scored the highest grade on a math test conveys sincere praise wrapped in an overtly critical form. Sarcasm, often regarded as a subset of irony, typically carries a more critical or mocking tone. It involves saying something ostensibly complimentary to convey disdain or contempt—e.g., “Oh, you’re so organized,” uttered in response to someone who forgot a crucial deadline. Both forms hinge on interpretive contrast: the audience must recognize a misalignment between what is said and what is meant. Crucially, decoding such utterances depends not only on lexical semantics and context but also on paralinguistic signals—particularly prosody [4]. Variations in intonation, pitch, and speech rhythm often signal the speaker’s true attitude and help listeners detect irony or sarcasm when the words alone might suggest sincerity [5–9].

Neuroscientific studies investigating irony and sarcasm in the auditory modality are scarce, but have highlighted the importance of regions involved in both language processing and social cognition, with the left inferior frontal gyrus (IFG) emerging as a consistent and central node. Matsui et al. [2] found that the left IFG pars orbitalis (IFGorb) is critically involved in integrating discourse context, prosodic information, and speaker intent during the comprehension of sarcastic utterances. Similarly, Obert et al. [10] reported activation of the bilateral IFG, along with the medial prefrontal cortex, during the processing of ironic speech. These regions contribute to both semantic evaluation and mentalizing processes, the ability to infer others’ mental states and communicative intentions. However, it remains unclear whether activation in the left IFG reflects the detection of prosodic–semantic incongruence [11, 12], the processing of irony or sarcasm, or a broader role in integrating prosody and semantics—an essential step in interpreting non-literal language—as suggested by Schirmer and Kotz [13].

To further elucidate how the brain integrates prosodic and semantic information during non-literal speech comprehension, Wittmann et al. [14] investigated the neural mechanisms underlying irony and sarcasm perception using naturalistic auditory dialogues. In this study, participants listened to brief conversational exchanges in which both the semantic context (established by a first speaker) and the prosody and semantics of the target utterance (from a second speaker) were systematically manipulated. This experimental design enabled the authors to investigate how various combinations of prosodic and semantic cues shaped participants’ evaluations of the target statements across tasks tapping into the key components of non-literal speech comprehension—namely, prosodic and semantic decoding, irony and sarcasm interpretation, and mentalizing. At the whole-brain level, non-literal speech elicited stronger activation than literal speech across the tasks in a broad network peaking in the left IFGorb, together with temporal regions associated with speech processing and areas classically involved in mentalizing, such as the medial prefrontal cortex and temporoparietal junction. To probe integration more directly, the authors then tested the prosody-by-semantics interaction within a set of regions of interest defined from this contrast, across the five tasks. This region-of-interest analysis revealed markedly heterogeneous integration profiles across regions and tasks, with the left IFGorb—the peak of the whole-brain contrast—showing the strongest and most consistent prosody-by-semantics interaction, present across all five tasks. However, its fractional factorial design—in which prosody and context factors covaried—and its restriction to a small number of a priori peak coordinates precluded strong inferences about which brain regions genuinely integrate prosodic and semantic information. To address this gap, the present study adopts a multivariate encoding approach that identifies such voxels directly, asking where the joint encoding of continuous prosodic and semantic features predicts neural responses beyond either modality on its own.

Encoding models have gained substantial attention in recent years for their ability to link linguistic representations with neural responses. In voxel-wise encoding models, features of interest—typically derived as embeddings from language stimuli—are used in regression analyses to predict how these features modulate BOLD responses in individual voxels. Early approaches relied on distributional semantic spaces, where each word was represented as a high-dimensional vector based on its normalized co-occurrence with a set of common English words across large text corpora, revealing a broad semantic network spanning temporal, parietal, and prefrontal cortex [15]. With the advent of deep learning, more powerful and context-sensitive models have emerged. For instance, Jain and Huth [16] demonstrated improved brain alignment using word embeddings derived from recurrent neural networks. More recently, transformer-based language models have enabled richer and more context-sensitive representations of linguistic input through attention mechanisms [17], capturing both syntactic and semantic dependencies over extended discourse. A growing body of research has demonstrated that embeddings derived from these models (e.g., GPT) can accurately predict neural responses across widespread language-related brain regions, with prediction performance increasing with layer depth and context span [18–22]. Beyond unimodal approaches, recent work has begun extending encoding models to jointly represent multiple input streams, for instance combining language and vision within a shared transformer-based framework [23]. While these studies highlight the power of deep language models in modeling language processing in the brain, they have primarily focused on textual or audio input alone. To our knowledge, no prior work has specifically investigated how the brain integrates semantic and prosodic information during language comprehension—a crucial step toward improving our understanding of social interactions, especially the neural basis of non-literal language such as irony and sarcasm.

In the present project, we aimed to investigate how the human brain integrates semantic and prosodic cues to comprehend non-literal language, such as irony and sarcasm. To this end, we derived custom contextualized semantic embeddings that model the relationship between an ironic statement and its preceding discourse through a context-attention mechanism, reflecting the fact that interpreting irony and sarcasm depends heavily on how context reshapes the meaning of a statement. In parallel, we extracted a standardized set of acoustic parameters that capture the affective prosody of the spoken statement. By combining these context-sensitive semantic representations with prosodic features, we fit voxel-wise ridge regression models to predict brain activity and identify regions whose responses are best explained by the integration of both semantic and prosodic information, rather than by either modality alone. We hypothesized that the left IFG functions as a central integration hub for prosodic and semantic information, alongside temporal speech regions [2, 13, 14]. To test whether such integration specifically depends on contextual weighting, we additionally compared these results against a statement-only baseline that encodes each utterance in isolation.

## Results

We analyzed the fMRI dataset of Wittmann et al. [14], here comprising 50 participants—the 45 participants analysed in that report together with 5 additional participants acquired under the same protocol—who listened to short two-character dialogues that could be interpreted literally or non-literally, depending on manipulations of the context semantics and of the target statement’s semantics and prosody. In the present project, we used data from the two tasks in which participants rated the perceived degree of irony and sarcasm. To identify brain regions integrating prosodic and semantic cues across these tasks, we trained voxel-wise linear ridge regression models across all participants and tasks, predicting fMRI activity from (1) semantic cues only, (2) prosodic cues only, or (3) both modalities combined. Semantic embeddings were obtained from the CamemBERTa-v2 model [24] and derived in two ways: a contextualized version transformed through a custom context-attention mechanism that weights each statement by its relevance to the preceding context—a key factor in non-literal speech comprehension—and a statement-only version that encodes each utterance in isolation. Each embedding type yields a text-only and a text+audio model, which together with the shared audio-only model give the five encoding models reported here; *text* and *text+audio* refer to the context-attention variant throughout unless the statement-only embeddings are named explicitly. Affective prosodic features of the statement were extracted with the extended Geneva Minimalistic Acoustic Parameter Set (eGeMAPSv02) [25]. All feature sets were entered alongside baseline covariates accounting for interindividual variability and task-related factors, helping the models isolate relevant information from the text and audio signals. For each model, we ran a 5-fold cross-validation and computed a brain score as the mean Pearson correlation between predicted and observed fMRI responses across test folds.

To identify voxels that genuinely integrate the two modalities, we required that each modality contribute predictive information that the other could not provide on its own. We therefore computed two conditional difference scores at each voxel: the unique contribution of audio over text, Δ*r*_audio|text_ = *r*_text+audio_ − *r*_text_, and the unique contribution of text over audio, Δ*r*_text|audio_ = *r*_text+audio_ − *r*_audio_. The significance of each conditional score was assessed with a permutation test in which only the added feature block was shuffled within each participant while the other block stayed aligned to the neural responses, so that the null isolates the variance uniquely attributable to the added modality (a Draper–Stoneman scheme; see Methods). A voxel was labelled *integrative* only if *both* conditional contributions were positive and significant—a minimum-statistic conjunction [26] whose displayed statistic is Δ*r*_int_ = min(Δ*r*_audio|text_, Δ*r*_text|audio_) and whose *p*-value is the larger of the two one-tailed conditional *p*-values. We computed this conjunction separately for each semantic embedding type, yielding one integration map based on the contextualized embeddings and one based on the statement-only embeddings. Voxel-wise *p*-values were corrected across in-brain voxels with the Benjamini–Hochberg false discovery rate (FDR) procedure at *q <* .05, and for reporting and visualization we retained positive, FDR-significant voxels within the top 1% of Δ*r*_int_ values, grouped into clusters of at least 10 contiguous voxels (26-connectivity); see S1 Fig for permutation diagnostics, and S2 Fig and S1 Table for the two underlying conditional-contribution maps. Only clusters in which every voxel met both criteria—surviving the Δ*r*_int_ threshold and showing FDR-corrected significance—were retained. Taken together, the combination of a high Δ*r*_int_ threshold, voxel-wise FDR correction, the requirement that both modalities contribute uniquely, and strict cluster-level requirements makes this analysis highly conservative, ensuring that the spatially contiguous clusters we observe reflect only the most robust and reliably coherent multimodal integration effects rather than scattered noise.

We first assessed the performance of the encoding models by computing voxel-wise brain scores reflecting the correlation between predicted and observed fMRI responses. The context-attention text model ranged from −0.087 to 0.142 (*M* = 0.013, *SD* = 0.019), with 75.4% of voxels showing positive scores, and the audio-only model from −0.072 to 0.132 (*M* = 0.012, *SD* = 0.018; 75.2% positive). The combined text+audio model performed slightly better, ranging from −0.071 to 0.142 (*M* = 0.015, *SD* = 0.018; 80.4% positive; S2 Table lists the brain scores of all models, and the whole-brain cluster tables are provided in S4 Table for text, S5 Table for audio, and S6 Table and S7 Table for the statement-only text and text+audio models). Turning to the unique contribution of each modality, both conditional scores were positive across roughly half of the brain but modest in magnitude: the unique audio contribution Δ*r*_audio|text_ averaged *M* = 0.002 (*SD* = 0.011; 51.3% of voxels positive) and the unique text contribution Δ*r*_text|audio_ averaged *M* = 0.003 (*SD* = 0.018; 55.3% positive). Because genuine integration requires both contributions jointly, the conjunction defining our integration maps is necessarily sparser and confined to a restricted set of regions, which we examine next. The text model predicted bilateral activity throughout the temporal cortices, with the strongest clusters extending across the superior and middle temporal gyri and the superior temporal sulcus (STG, MTG, STS) in both hemispheres, together with smaller orbitofrontal and superior frontal clusters (Fig 1A; S4 Table). The audio model recovered a more scattered, right-dominant temporal profile: its most extensive cluster peaked in the right STG and extended across adjacent superior and middle temporal regions (STG, STS, MTG), consistent with the expected cortical topography of prosodic processing. Prosodic features additionally predicted orbitofrontal responses. The map also comprised numerous smaller clusters distributed across occipital, parietal, and cerebellar regions (Fig 1B; S5 Table). The combined text+audio model reproduced this bilateral temporal pattern (S3 Fig; S8 Table). Beyond confirming that each model recovers its expected cortical topography, these maps localize, within a non-literal speech context, the semantic and prosodic representations on which the subsequent integration analysis builds.

**Fig 1.**
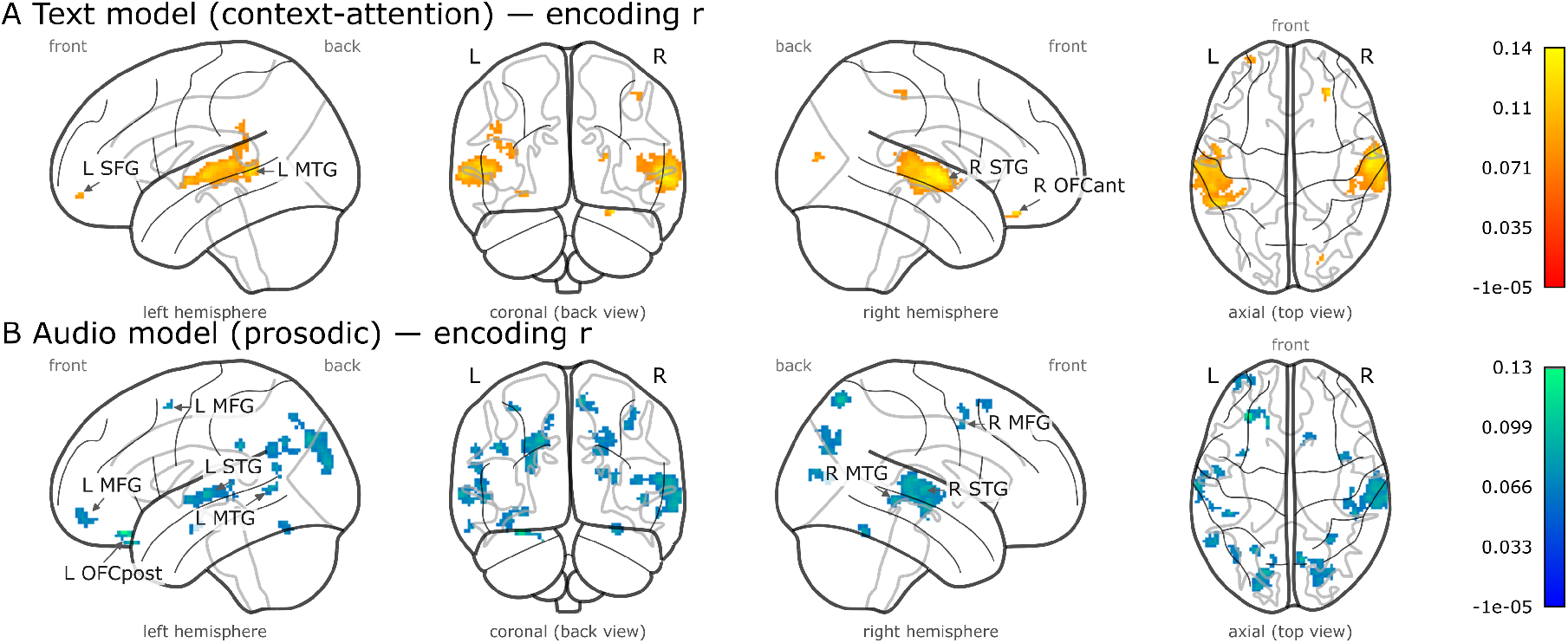
Unimodal encoding performance for the semantic and prosodic models. Glass-brain maps of voxels surviving the top 1% positive threshold (FDR-corrected *q <* .05, clusters ≥ 10 contiguous voxels). Panel **A** shows brain encoding performance (Pearson *r*) for the text model (context-attention embeddings) and Panel **B** for the audio (prosodic) model. Each panel is scaled to its own maximum, as indicated by the accompanying colour bars. Labelled clusters are the strongest temporal and frontal peaks per model. MFG: middle frontal gyrus; SFG: superior frontal gyrus; MTG: middle temporal gyrus; STG: superior temporal gyrus; OFCant: anterior orbital gyrus; OFCpost: posterior orbital gyrus.

Restricting the analysis to voxels where both modalities contributed unique variance, the context-attention model localized integration to a small set of regions beyond auditory cortex (Fig 2A; Table 1A). The strongest peak fell in the left IFGorb, together with a left precentral cluster, the right superior temporal gyrus, and a right paracentral cluster. Because this conjunction is directional and conservative, the surviving clusters are spatially focal; their modest extent, relative to the unimodal encoding maps, is expected for an effect that requires both modalities to contribute jointly.

**Table 1.** Integration clusters (minimum-statistic conjunction of the two conditional contributions) for the context-attention and statement-only embeddings. Each panel lists the brain clusters in which *both* the unique audio contribution (Δ*r*_audio|text_) and the unique text contribution (Δ*r*_text|audio_) were positive and FDR-significant (*q <* .05), within the top 1% of Δ*r*_int_ = min(Δ*r*_audio|text_, Δ*r*_text|audio_) and forming clusters of at least 10 contiguous voxels. Panel **A** corresponds to the context-attention variant and Panel **B** to the statement-only variant. For each cluster we report its size (in voxels), mean and peak Δ*r*_int_, FDR-corrected *q*-value at the peak voxel, peak and centre-of-mass (CoM) MNI coordinates, anatomical region (AAL atlas), and hemispheric laterality. Within each panel, clusters are ranked in descending order of peak Δ*r*_int_.

| # | Size<br>(vox) | Mean<br>$\Delta r_{\text{int}}$ | Peak<br>$\Delta r_{\text{int}}$ | Peak $q$<br>FDR | Peak MNI Coord | CoM MNI Coord | Region | Lat |
| --- | --- | --- | --- | --- | --- | --- | --- | --- |
| **A. Context-attention** |  |  |  |  |  |  |  |  |
| 1 | 14 | 0.039 | 0.058 | .034 | (−28, 38, −2) | (−32.3, 37.1, −4.6) | IFGorb | L |
| 2 | 10 | 0.037 | 0.045 | .034 | (50, −32, 14) | (46.6, −33.2, 16.4) | STG | R |
| 3 | 10 | 0.037 | 0.044 | .034 | (−42, −2, 44) | (−42.2, −1.6, 46.0) | PreCG | L |
| 4 | 15 | 0.039 | 0.044 | .034 | (2, −38, 68) | (2.3, −36.5, 68.1) | PCL | R |
| **B. Statement-only** |  |  |  |  |  |  |  |  |
| 1 | 117 | 0.036 | 0.084 | .037 | (−64, −26, 4) | (−60.7, −23.1, 3.1) | STG | L |
| 2 | 82 | 0.034 | 0.056 | .037 | (60, −8, 0) | (61.4, −8.2, −0.8) | STG | R |

**Note.** IFGorb: inferior frontal gyrus, orbital part; STG: superior temporal gyrus; PreCG: precentral gyrus; PCL: paracentral lobule; CoM: centre of mass; Lat: laterality; L: left; R: right.

**Fig 2.**
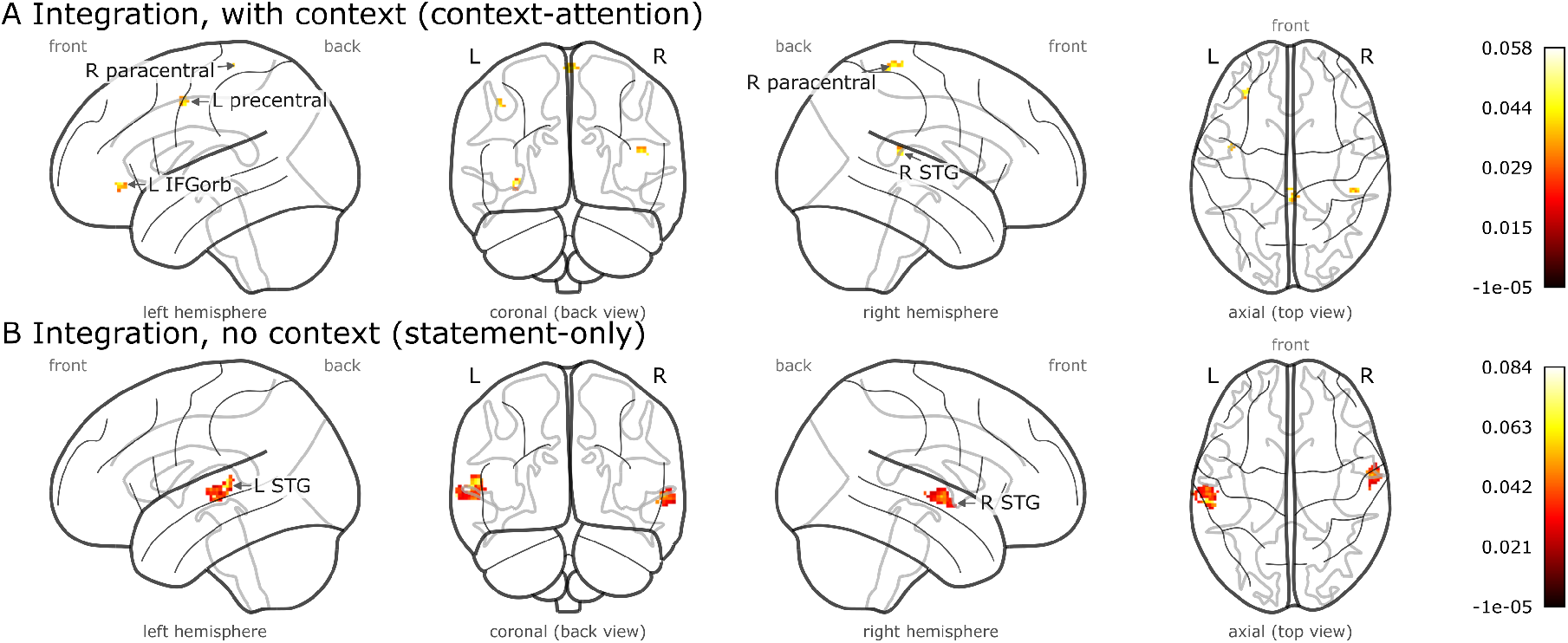
Multimodal integration of prosodic and semantic cues depends on contextual weighting. Glass-brain maps of the integration conjunction Δ*r*_int_ = min(Δ*r*_audio|text_, Δ*r*_text|audio_), retaining voxels where *both* modalities contributed unique predictive variance (positive and FDR-significant at *q <* .05, top 1%, clusters ≥ 10 contiguous voxels). Panel **A** (context-attention embeddings): integration engages the left inferior frontal gyrus pars orbitalis (IFGorb) alongside the right superior temporal gyrus. Panel **B** (statement-only embeddings): integration is confined to bilateral temporal cortex. Each panel is scaled to its own maximum, as indicated by the accompanying colour bars; the contrast between them concerns *where* integration arises rather than its magnitude. The paracentral cluster straddles the midline and is therefore visible in both sagittal projections. IFGorb: inferior frontal gyrus, orbital part; STG: superior temporal gyrus.

To test whether this frontal recruitment depended on modeling discourse context, we repeated the conjunction with statement-only embeddings that encode each utterance in isolation. The statement-only combined model performed comparably to the context-attention model (from −0.061 to 0.129; *M* = 0.015, *SD* = 0.017; 80.8% positive; S4 Fig), yet its integration profile was markedly different: integration was confined to bilateral temporal cortex, with a large cluster spanning the left superior and middle temporal gyri and one in the right superior temporal gyrus (Fig 2B; Table 1B); the two underlying conditional contributions for the statement-only embeddings are mapped in S5 Fig and listed in S3 Table. Critically, the left IFGorb did not survive the reporting threshold under the statement-only embeddings. Under FDR correction alone it remained the strongest of 49 integration clusters with context-attention embeddings but ranked 33rd of 49 with statement-only embeddings (S9 Table, S10 Table). Thus contextual weighting shifted the left IFGorb from a marginal to the dominant integration site, whereas temporal integration was present regardless.

## Discussion

In this study, we combined voxel-wise encoding models with a context-sensitive semantic representation and a conjunction-based integration test to identify where the brain combines prosodic and semantic cues during the comprehension of non-literal speech, such as irony and sarcasm. By requiring that a voxel be better predicted by the joint model than by *either* modality alone, we isolated a small set of regions in which the two information streams are genuinely integrated rather than merely co-represented. The anatomy of this set depended critically on whether the semantic representation incorporated discourse context: integration was largely confined to temporal speech cortex when statements were encoded in isolation, but extended to the left IFGorb when each statement was weighted by its relevance to the preceding context. This dissociation suggests that the left IFGorb is recruited specifically when comprehension requires combining semantic, prosodic, and contextual information at once—the defining demand of ironic and sarcastic speech.

When statements were encoded in isolation, the regions integrating prosody and semantics were dominated by the bilateral temporal cortex, peaking in the left and right superior temporal gyri, with the left cluster extending into the middle temporal gyrus. These regions lie at the core of the speech-processing hierarchy: the left temporal cortex has classically been associated with lexical–semantic analysis [27, 28], and the right temporal cortex with the decoding of emotional prosody [29–31], although this hemispheric division is far from strict [13, 32]. That the strongest integration in the statement-only model falls in these regions suggests that, in the absence of contextual weighting, prosodic and semantic information are combined primarily at the level of auditory and speech representations themselves, rather than in downstream regions.

The right superior temporal cluster, which also survived the conjunction under contextual weighting, is consistent with proposals that the posterior superior temporal cortex acts as a convergence zone binding auditory-object representations with affective multimodal cues [33, 34]; our findings extend this role to the integration of affective prosody with semantics within the auditory domain.

Modeling discourse context changed this picture, with the left IFGorb emerging as the strongest integration cluster—a region that fell far below the reporting threshold under the statement-only embeddings. The left IFGorb has repeatedly been implicated in non-literal language: Matsui et al. [2] identified it as critical for integrating discourse context, prosody, and speaker intent during sarcasm comprehension, and it sits at the interface of semantic and emotional—including prosodic—processing [13, 35]. Most directly, Wittmann et al. [14] found the left IFGorb to be the peak of non-literal speech processing across tasks and, in a region-of-interest analysis, reported a significant prosody-by-semantics interaction there in every task, including the irony and sarcasm tasks. That interaction, however, was defined over categorical valence conditions in which prosody and context covaried, so it could not separate prosody–semantics integration from a context–semantics interaction, nor establish whether the region genuinely encodes the combination of prosodic and semantic information rather than the other processes that distinguish non-literal from literal speech. Our results help disentangle this: the left IFGorb emerges as the dominant integration site specifically when semantics are weighted by context, suggesting that it integrates prosody not with the literal content of an utterance but with its context-dependent, pragmatically reshaped meaning. This shift is not an artefact of our reporting threshold: under FDR correction alone the left IFGorb was the strongest of 49 integration clusters with context-attention embeddings (29 voxels; Δ*r*_int_: mean 0.021, peak 0.058) and ranked 33rd of 49 with statement-only embeddings (16 voxels; Δ*r*_int_: mean 0.003, peak 0.012; S9 Table, S10 Table). This positions the left IFGorb as a higher-order convergence region combining context, semantics, and prosody, one step beyond the temporal integration observed for isolated statements. In other words, the IFGorb appears to be engaged for integration specifically when it cannot be resolved within auditory cortex alone—when prosody must be combined with a meaning that has itself been reshaped by context, as is constitutive of irony and sarcasm.

Beyond the IFGorb, prosodic features also predicted orbitofrontal responses, which fits the proposal that an orbitofrontal network, connected to the amygdalae and the temporal voice areas, infers contextual value from the relation between context and prosody—the very demand irony and sarcasm impose [34]. This role is consistent with the region’s place at the evaluative stage of prosody comprehension [36], and with its recruitment when semantic and prosodic cues conflict [37]. Other regions in which prosody and semantics were integrated, or in which prosody contributed unique predictive value beyond semantics, fell in motor and somatosensory cortices. The integration conjunction retained a left precentral and a right paracentral cluster—the latter lying on the medial wall, where the primary motor and somatosensory strips converge—and the unique contribution of prosody over semantics peaked in the right postcentral gyrus, additionally engaging the left supplementary motor area and the right precentral gyrus (S1 Table). This is reminiscent of the network described by Skipper et al. [38], who found that motor and somatosensory regions were reliably recruited during audiovisual speech perception, but notably not during audio-only or visual-only speech. Could it be that, just as motor and somatosensory areas support the integration of auditory and articulatory information in face-to-face speech, they also subserve a broader integrative role in binding the prosodic and semantic dimensions of spoken language? This possibility would align with an embodied perspective, in which experience with producing and perceiving speech not only links sounds to motor patterns but also anchors prosodic variation in articulatory and bodily experience, so that speech is understood not simply as an auditory signal but as an embodied act [39], a view reinforced by the recruitment of these same motor and somatosensory regions during emotional prosody production [40]. More broadly, sensorimotor circuits have been proposed to form a cortical basis for language, actively shaping comprehension rather than merely reflecting it [41], and even passive listening to speech recruits premotor regions that overlap those engaged during its production [42]. The way prosody and semantics are dynamically integrated through sensorimotor grounding could then shape how listeners distinguish and interpret meaning, including in non-literal forms of speech. Although this interpretation remains highly speculative, future work should investigate directly whether these regions contribute to binding prosody and semantics rather than merely tracking articulatory information. Further, similar paradigms could examine how multiple modalities, including visual cues, semantics, and prosody, are integrated during speech comprehension.

Our study has several limitations. First, the brain scores obtained were modest, with the highest value across all models reaching *r* = 0.151 (statement-only text model; S2 Table). For comparison, Caucheteux et al. [22] reported a maximum correlation of *r* = 0.23. Although our correlations were relatively low, the conditional Δ*r* values demonstrated that each modality contributed unique predictive variance, and the conjunction test isolated voxels where the two were jointly integrated. Because this conjunction is conservative—requiring both contributions to be independently significant—the resulting integration clusters were small, and their spatial extent should be interpreted with corresponding caution. Relatedly, the top 1% percentile threshold and the minimum cluster-size criterion are pragmatic rather than principled choices: because a large number of voxels reached FDR significance in both the unimodal and the integration analyses, we deliberately restricted reporting to the voxels showing the strongest effects and to spatially coherent clusters, both to isolate the most robust integration and for clarity of visualization. Different thresholds would yield somewhat more or less extensive maps, and the specific spatial extents should therefore be read as descriptive rather than definitive. The full FDR-corrected cluster tables are provided in S9 Table and S10 Table, and the principal dissociation does not depend on the percentile criterion. Several factors may account for the modest absolute predictivity. Notably, given the limited number of trials available per participant to train the encoding models, we concatenated data across all participants to ensure sufficient statistical power. This procedure may have introduced variability, as inter-individual differences in baseline activity, neuroanatomy, and cognitive processing could have reduced the model’s sensitivity to the representational features derived from text embeddings and audio inputs; we included individual covariates in every model to partly offset this. A second limitation concerns the temporal resolution of our analysis. Because our focus was on the overall meaning of the ironic statements rather than word-level processing at each repetition time (TR), we used mean activation spanning from stimulus onset to the end of the evaluation. While this approach allowed us to capture holistic processing, it may not align with methods used in prior work that modeled embeddings at the TR level and employed finite impulse response functions to characterize the delayed dynamics of the BOLD signal [15, 21, 22]. To mitigate this issue, we applied a weighted mean with hemodynamic response function (HRF) weights, assigning greater importance to TRs corresponding to the expected peak of neural activity. Nevertheless, while necessary for our primary objective of capturing the integrated meaning of the full ironic statement, this strategy remains an approximation and may not fully capture the fine-grained temporal alignment between embeddings and neural responses.

Beyond these limitations, we believe that our work makes several valuable contributions. Methodologically, we introduce a principled approach to localize multimodal integration: we test the unique contribution of each modality with a block-restricted (Draper–Stoneman) permutation scheme and retain only voxels where both contributions are jointly significant, using a minimum-statistic conjunction [26]. This isolates genuine integration from the mere co-representation of two correlated streams and could be extended beyond two modalities to characterize integration hubs across the brain. In addition, whereas previous studies leveraged forecast windows by concatenating embeddings from fixed spans of past or future context (e.g., Caucheteux et al. [22]), we implemented an attention-based mechanism that selectively weights contextual information relative to the target sentence (or word). This approach allowed us to model context-sensitive meaning more flexibly and more closely reflecting human pragmatic processing, where certain contextual cues are weighted more heavily than others. Such selective integration is particularly relevant for irony and sarcasm, which rely on nuanced contrasts between local linguistic input and broader discourse context. Taken together, these innovations provide a foundation for future work to refine methodological choices and further probe the neural basis of multimodal integration. More broadly, they also contribute to the development of computational approaches for studying pragmatic aspects of language processing, such as non-literal speech.

## Conclusion

In this study, we examined how the brain integrates prosodic and semantic cues during the comprehension of non-literal speech using voxel-wise encoding models and a conjunction-based test of multimodal integration. Requiring that each modality contribute predictive information beyond the other localized integration to a small, anatomically interpretable set of regions whose composition depended on contextual weighting. When statements were encoded in isolation, integration was confined to the bilateral temporal speech cortex; when each statement was weighted by its relevance to the preceding context, integration additionally engaged the left inferior frontal gyrus pars orbitalis. This dissociation indicates that the left IFGorb becomes the dominant site of prosody–semantics integration in ironic and sarcastic speech only when meaning has been shaped by context, and it clarifies the integrative role of this region that earlier univariate work could identify but not interpret. The methodological framework introduced here—combining language-model-derived, context-sensitive representations with conditional, conjunction-based integration tests—provides a flexible tool for probing where and under what conditions the brain combines information sources during naturalistic communication, and extends the questions that can be asked of naturalistic neuroimaging data beyond those addressable with condition-based designs. Looking ahead, future work should investigate the temporal dynamics of these integration processes and broaden the range of modalities and contextual variables incorporated into predictive models.

## Materials and methods

### Participants

Our dataset included 50 participants (26 males and 24 females, *M*_age_ = 23.18, *SD*_age_ = 3.62) recruited at the University of Geneva, Switzerland: the 45 participants analysed by Wittmann et al. [14] and 5 additional participants acquired under the same protocol and ethics approval. All participants provided written informed consent on a signed consent form before taking part and received monetary compensation, and the study adhered to the Declaration of Helsinki and was accepted by the local Ethics committee for research (CCER; approval number 2021-01219). Eligibility criteria required participants to be between 18 and 45 years old, to have French as their native language, not to be pregnant, not to suffer from claustrophobia, and to have no known hearing or speech impairments and no history of psychiatric disorders. Recruitment and data collection ran from 22 June 2023 to 13 March 2024.

### Stimuli and tasks

Our stimuli consisted of recorded sentences in French featuring short dialogues between two characters. Specifically, the first character delivered an utterance, referred to as the context (*M*_duration_ = 2.10, *SD*_duration_ = 0.33), and the second character responded with the target statement (i.e., the one that could be ironic or not; *M*_duration_ = 1.06, *SD*_duration_ = 0.21). We created 16 scenarios in which the semantics of the context and the semantics and prosody of the statement were manipulated. For a given scenario, the context could be stated either positively or negatively (e.g., “my husband won a lot of money in the lottery” vs. “this player lost a lot of money in the lottery”) with a monotone prosody, while the target statement could be stated either positively or negatively (e.g., “he is lucky” vs. “he is unlucky”) and delivered with a positive or negative prosody. In the fMRI study, only four conditions were included: sarcastic irony (negative context, positive statement, negative prosody), praise irony (positive context, negative statement, positive prosody), negative sincerity (negative context, negative statement, negative prosody), and positive sincerity (positive context, positive statement, positive prosody). In the current project, we only analyzed the data from the irony and sarcasm tasks in which participants were required to evaluate the degree of perceived irony from *not ironic at all* (1) to *very ironic* (5) and the degree of perceived sarcasm from *not sarcastic at all* (1) to *very sarcastic* (5). The experimental design is summarized in Fig 3. For further details about the experimental design, procedure, and image acquisition, see Wittmann et al. [14].

**Fig 3.**
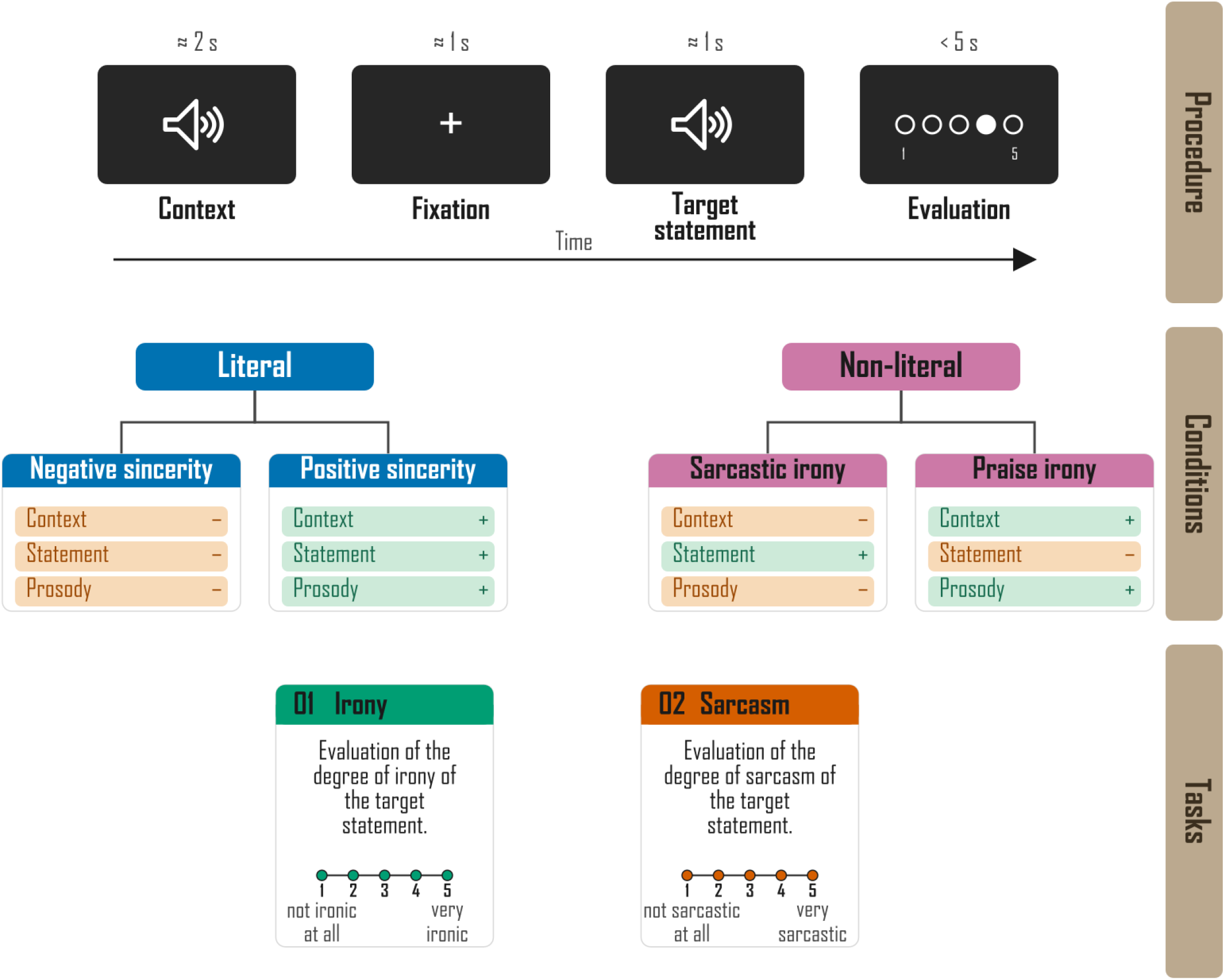
Summary of the fMRI experimental plan. **Top:** trial procedure. Each trial presented the spoken context (≈2 s), a fixation cross (≈1 s), the spoken target statement (≈1 s), and a self-paced evaluation on a five-point scale (*<*5 s). **Middle:** the four conditions, defined by the valence of the context, of the statement semantics, and of the statement prosody. Literal conditions align all three cues (negative sincerity, positive sincerity), whereas non-literal conditions oppose statement semantics to context and prosody (sarcastic irony, praise irony). **Bottom:** the two tasks analysed here, in which participants rated the perceived degree of irony and of sarcasm of the target statement. Adapted from Wittmann et al. [14].

### Data preprocessing

Preprocessing of functional images was performed using SPM12 (Wellcome Trust Centre for Neuroimaging, London, UK). All volumes were realigned to the first image to correct for motion, coregistered, normalized to Montreal Neurological Institute (MNI) space [43], and spatially smoothed with a 6 mm full-width at half-maximum Gaussian kernel. After standard preprocessing, nuisance signals were removed via linear regression. Confound regressors included the six rigid-body motion parameters estimated during realignment (three translations, three rotations), their temporal derivatives, and the squares of both (24 motion regressors total). In addition, mean signals from white matter and cerebrospinal fluid were extracted using participant-specific tissue masks thresholded at a probability of 0.9 and resampled to functional resolution, yielding two physiological noise regressors. A high-pass filter with a cutoff of 0.01 Hz was applied simultaneously to remove slow signal drifts. Each voxel’s residual time series was then z-scored by subtracting its temporal mean and dividing by its standard deviation, separately for each participant and run, to account for differences in baseline signal across individuals and sessions. Because our analyses focused on the neural representation of the entire ironic utterance—whose interpretation requires integrating semantic and prosodic cues at the sentence level—we needed a single trial-level neural estimate rather than TR-wise or word-level responses. To obtain a summary measure aligned with this integrated representation, we computed an HRF-weighted mean of the BOLD signal. For each trial, we extracted the BOLD time series spanning both the listening period and the subsequent evaluation period, since the hemodynamic response to short auditory events typically peaks during this later window and cognitive resolution of irony often continues into evaluation. Each time point was weighted by the canonical Glover HRF convolved with the statement onset and duration, and only positive HRF values were retained, yielding a single value per voxel per trial that reflects the full neural response with improved temporal specificity. Finally, all resulting trial-level brain volumes were concatenated across participants and runs and re-normalized to zero mean and unit variance, ensuring consistent scaling for subsequent analyses.

### Semantic embeddings

Semantic features for each target statement were derived from embeddings obtained from the final layer of CamemBERTa-v2 [24], a 12-layer transformer model implemented through the Hugging Face Transformers library [44]. Each token embedding produced by the model is a 768-dimensional vector. We derived semantic features in two complementary ways. The first, a *statement-only* embedding, encodes each target utterance in isolation, without reference to the preceding discourse. The second, a *context-attention* embedding, models how the surrounding context modulates the interpretation of the target statement through a custom attention-based mechanism.

Because interpreting a target statement—particularly when it is ironic or sarcastic—depends strongly on the preceding discourse, this second variant was designed to capture how contextual semantics reshape the representation of the target utterance. Comparing the two embedding types allows us to assess how incorporating context affects the neural encoding of non-literal meaning.

#### Statement-only embeddings

For the statement-only variant, we encoded the target statement and computed its mean-pooled embedding:

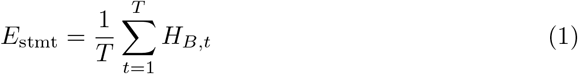

where *E*_stmt_ ∈ ℝ^1*×d*^ is the statement embedding, *H*_*B,t*_ ∈ ℝ^*d*^ are the token-level hidden states from the final transformer layer, and *T* is the length of the target-statement token sequence. This yields a 768-dimensional feature vector encoding the semantic content of the statement independently of context.

#### Context embeddings

For the context-attention variant, we first encoded the context and computed its mean-pooled embedding:

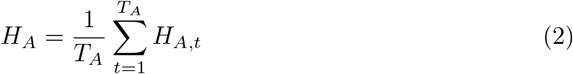

where *H*_*A*_ ∈ ℝ^1*×d*^ is the context embedding and *H*_*A,t*_ ∈ ℝ^*d*^ are token-level hidden states from the final transformer layer. *T*_*A*_ is the length of the context token sequence.

#### Attention over the target statement

We then encoded the target statement to obtain its token-level embeddings *H*_*B,t*_ ∈ℝ^*d*^, forming the matrix:

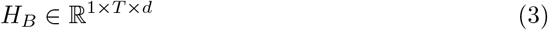

To model the influence of context on individual tokens in the target statement, we computed attention weights by applying a softmax function to the dot-product similarity between each target token and the context embedding:

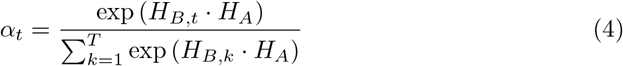

yielding *α* ∈ ℝ^1*×T*^. These weights assign greater importance to target-statement tokens that are more semantically aligned with the context.

#### Context-weighted target embeddings

Using the attention weights, we derived a context-weighted embedding of the target statement:

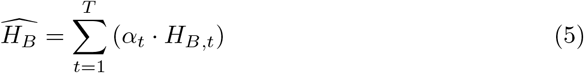

where 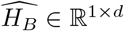. The operator · denotes the scalar multiplication of the vector *H*_*B,t*_ by the weight *α*_*t*_.

#### Context–statement incongruence

To capture semantic incongruity—an essential feature of irony and sarcasm—we computed the element-wise difference between the context-weighted target representation and the context embedding:

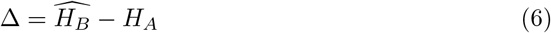

with Δ ∈ ℝ^1*×d*^. This vector quantifies the discrepancy between the context-weighted representation of the target statement and the context embedding.

#### Final combined embeddings

For the context-attention variant, we concatenated the context-weighted representation and the incongruence vector to obtain the final semantic embedding:

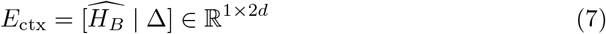

resulting in a 1536-dimensional feature vector for each context–statement pair, encoding both the semantic content of the statement in relation to the context and its deviation from contextual expectations (Fig 4). Both the statement-only (*E*_stmt_) and context-attention (*E*_ctx_) feature vectors were subsequently *z*-scored across items to ensure comparability, and each was entered into a separate set of encoding models.

**Fig 4.**
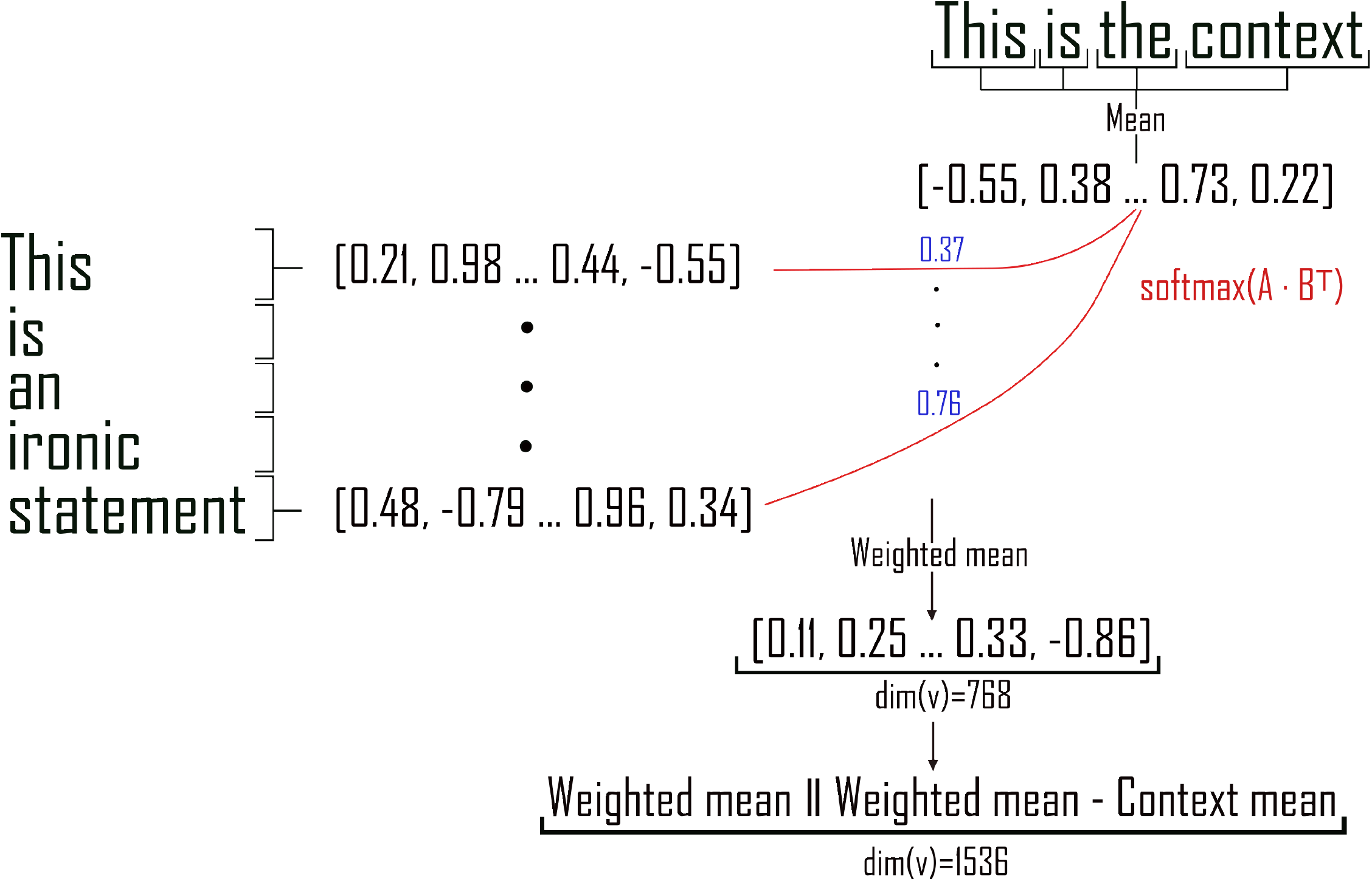
Custom context-weighted embeddings capturing statement meaning and contextual deviation. The diagram illustrates the computation of context-weighted semantic embeddings for ironic statements. The context sentence (top) is encoded and mean-pooled to produce a context embedding. The target statement (left) is tokenized and encoded to obtain token-level embeddings. Attention weights are computed using softmax over the dot-product similarity between each target token and the context embedding, shown as the red curve with example weights. These weights are used to compute a context-weighted mean of the target statement tokens. The final semantic embedding concatenates the context-weighted representation with the context–statement incongruence vector, resulting in a 1536-dimensional feature vector encoding both the semantic content of the statement relative to context and its deviation from contextual expectations.

### Prosodic features

To extract prosodic features from the target statement audio recordings, we used the openSMILE toolkit [45] with the eGeMAPSv02 feature set [25] at the functionals level. This feature set is specifically designed to optimally capture affective prosody through 88 parameters—a carefully selected set of low-level descriptors together with their statistical functionals—covering acoustic dimensions such as pitch, loudness, spectral balance, and voice quality. For each audio file in the dataset, we computed these features, resulting in a fixed-length 88-dimensional vector that summarizes the global prosodic characteristics of the recording. To ensure comparability across recordings, the resulting prosodic feature vectors were then *z*-scored.

### Baseline features

Because the dataset included multiple tasks, participants, and experimental conditions, we incorporated a set of baseline covariates into each regression model to help account for these sources of variability and to guide the model toward relevant information within the text and audio features. These baseline features included participant number, gender, age, task type, evaluation score, context semantics, statement semantics, and statement prosody. Categorical variables were encoded using one-hot encoding with a reference category to avoid multicollinearity. For the combined text+audio models, where both factors are present, we additionally included the interaction between statement semantics and statement prosody as a covariate (the products of their one-hot indicators), so that any variance attributable to the categorical semantics–prosody pairing was captured by the baseline rather than being absorbed by the text and audio feature blocks; this interaction was omitted from the unimodal models, in which only one of the two factors exists. Missing evaluation scores were imputed using the median, and evaluation values were subsequently 0–1 normalized using min–max scaling, preserving the rank ordering of observations while avoiding assumptions about equal spacing between values. Age was standardized using z-score normalization.

### Regression models and hyperparameter optimization

In each voxel-wise linear ridge regression model, the ridge regularization parameter *α* was optimized using a leave-one-participant-out cross-validation scheme, in which the data from one participant served as the test fold while the model was trained on data from all other participants. This procedure was performed separately for each voxel and each model. Prior to regression, a singular value decomposition of the training stimulus matrix was computed, and singular values below 10^−10^ were discarded to remove noise components and improve numerical stability. A final 5-fold cross-validation was then performed for each model, ensuring that data from the same participant were kept within the same fold and that participant splits were consistent across all models.

### Brain scores

To evaluate how well each voxel’s activity was predicted by our feature sets, we computed a brain score for each participant and voxel. The brain score is a model-to-brain predictivity metric that quantifies how well a linear encoding of model features predicts neural responses; it was originally developed to compare artificial neural networks with visual cortex [46], later formalized as the Brain-Score benchmark [47], and adapted to language models by Caucheteux et al. [21, 22], whose approach we follow here. Specifically, for each model—semantic-only, prosodic-only, and their combination—we fitted a voxel-wise linear ridge regression to predict the fMRI time series from the feature embeddings. This linear mapping was estimated on training data and evaluated using a 5-fold cross-validation. The brain score for each voxel was defined as the average Pearson correlation between the predicted and actual fMRI responses across the test folds. This yielded three whole-brain correlation maps, one for each model.

To assess whether a voxel’s activity was better explained by the integration of semantic and prosodic cues than by either modality alone, we quantified the unique contribution of each modality over the other. For each voxel we computed two conditional difference scores:

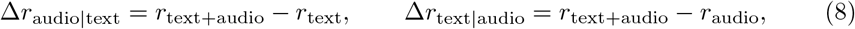

where *r*_text+audio_ is the brain score of the combined model and *r*_text_ and *r*_audio_ are the brain scores of the unimodal models. A positive Δ*r*_audio|text_ indicates that adding the prosodic features improves prediction beyond the semantic features alone, and conversely for Δ*r*_text|audio_. A voxel was considered to integrate the two modalities only if both conditional contributions were positive and statistically significant; the integration statistic reported is the minimum-statistic conjunction Δ*r*_int_ = min(Δ*r*_audio|text_, Δ*r*_text|audio_), with conjunction *p*-value equal to the larger of the two conditional *p*-values [26].

We computed this conjunction separately for the context-attention and the statement-only embeddings, yielding one integration map for each and allowing us to test whether integration in a given region depended on contextual weighting.

### Permutation, significance, and spatial filtering

To obtain voxel-wise significance estimates, we ran non-parametric permutation tests (1,000 iterations). The two conditional contributions were tested with a Draper–Stoneman scheme [48, 49] in which, within each participant, only the *added* feature block was permuted across trials while the other block and all nuisance regressors stayed aligned to the neural responses: for Δ*r*_audio|text_ the audio block was shuffled and the text block kept fixed, and vice versa for Δ*r*_text|audio_. The text block comprised the context-weighted embeddings and categorical semantic features, and the audio block the openSMILE-derived prosodic embeddings and prosodic base features. Because the combined model also included the semantics-by-prosody interaction described above—the products of the categorical one-hot indicators, whose parents sit in the text and audio blocks—these terms were not permuted independently but regenerated from the shuffled parent block on every iteration, so that the interaction always followed the permuted design. This isolates the variance uniquely attributable to the added modality while preserving temporal autocorrelation, the aligned modality, and all baseline covariates. Permuting the feature block of interest directly removes its association not only with the neural responses, as intended, but also with the other regressors retained in the model—the aligned modality and the categorical design variables. The direction of any resulting bias therefore depends on the correlation between the permuted block and those retained regressors, and can in principle be either liberal or conservative [48, 49]. The residualization- and orthogonalization-based corrections that best address this (the Freedman–Lane and Smith/Dekker schemes) are formulated for ordinary least squares and do not transfer directly to a ridge setting; we therefore kept the categorical design variables as baseline regressors in every model and report the resulting permutation diagnostics in S1 Fig. These show that the unimodal and combined encoding-model nulls were well-behaved—centred near zero with approximately uniform *p*-value distributions—whereas the nulls of the conditional contributions were more concentrated; because voxel-wise conditional significance is consequently liberal on its own, integration was never inferred from a single conditional map, but only from the conjunction of *both* contributions together with the top 1% Δ*r*_int_ and cluster-extent criteria.

For each permutation we refit the relevant ridge models with the same voxel-wise hyperparameters as in the observed analysis and recomputed the corresponding score. Voxel-wise one-tailed *p*-values were obtained as

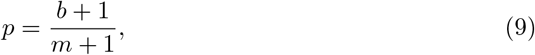

where *b* is the number of permutations yielding a score at least as large as the observed one and *m* = 1,000, an estimator that avoids zero *p*-values [50]. For the integration conjunction, the voxel *p*-value was the larger of the two conditional one-tailed values [26]. All *p*-values were corrected across in-brain voxels using the Benjamini–Hochberg FDR at *q <* .05.

To isolate spatially coherent effects, we retained voxels that were simultaneously positive, FDR-significant (*q <* .05), and within the top 1% of positive Δ*r* values, and grouped them into clusters using 26-connectivity (voxels sharing a face, edge, or corner), keeping only clusters of at least 10 voxels. This suppresses isolated single-voxel responses while preserving anatomically focal effects.

## Data and code availability

The behavioral and fMRI data analyzed here are openly available in the Yareta repository at https://doi.org/10.26037/yareta:cjiya654hzhivo5fotfygs5quu. All analyses were performed using Python. The ridge regression analyses were performed using a code adapted from Lebel et al. [51] available on GitHub at https://github.com/HuthLab/deep-fMRI-dataset. The text embeddings were extracted using the transformers library [44] and audio features extracted using the openSMILE library [45]. Our full project code is available at https://github.com/AdrienWitt/Prosody_Semantics_Integration_Irony_Encoding and permanently archived at https://doi.org/10.5281/zenodo.21411530.

## Acknowledgments

We thank Olivier Renaud (University of Geneva) for discussion of the permutation scheme used to test the conditional contributions.

## Supporting information

**S1 Fig.**
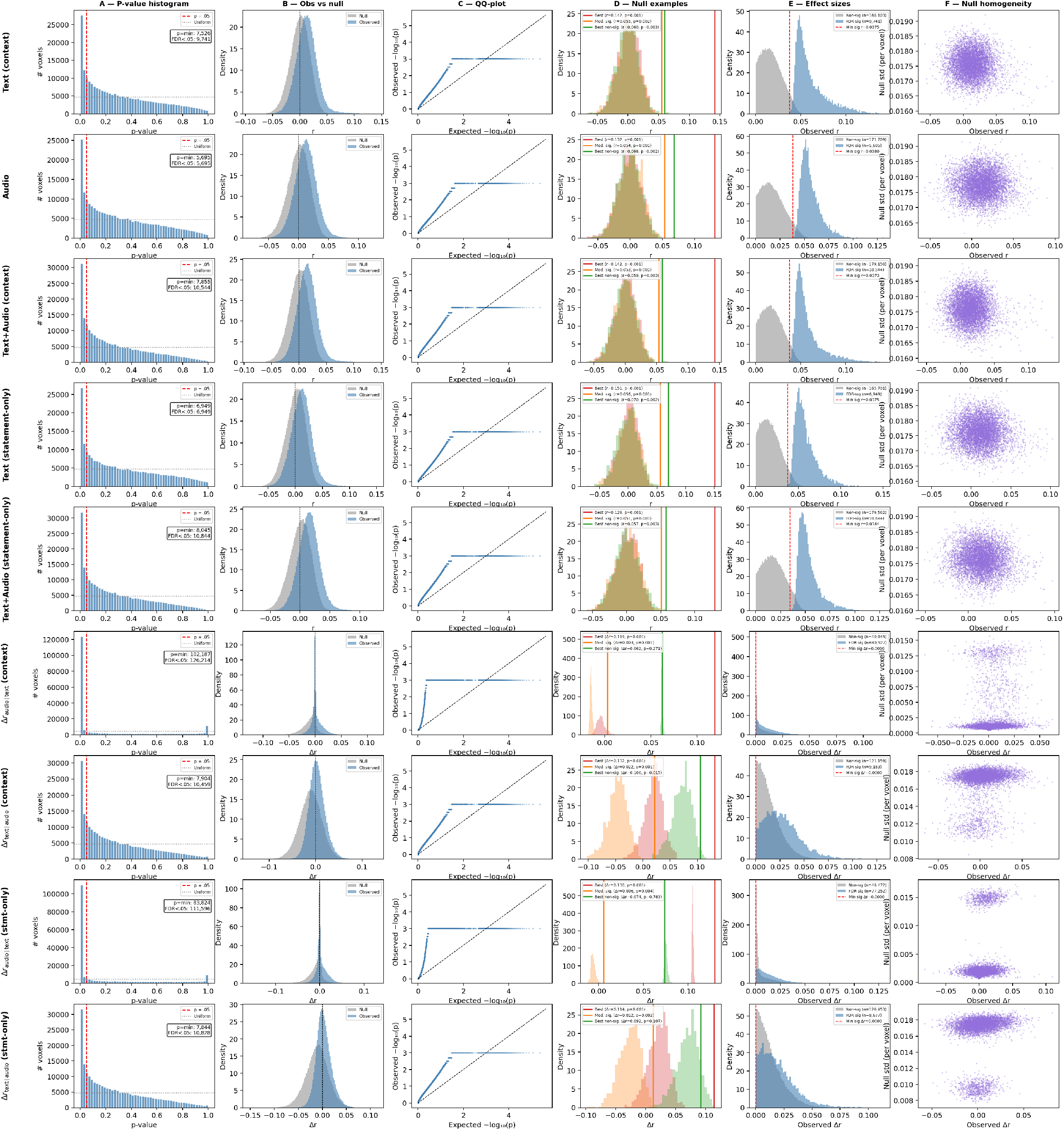
Permutation diagnostics. Distribution of permuted versus observed Δ*r* statistics and related diagnostics for the five encoding models and the four conditional contributions. Each row corresponds to one model; columns show (A) the *p*-value histogram, (B) observed versus null score distributions, (C) the QQ-plot of observed against expected *p*-values, (D) example null distributions with the best significant and best non-significant voxels, (E) observed effect sizes split by significance, and (F) null homogeneity across voxels.

**S1 Table.**
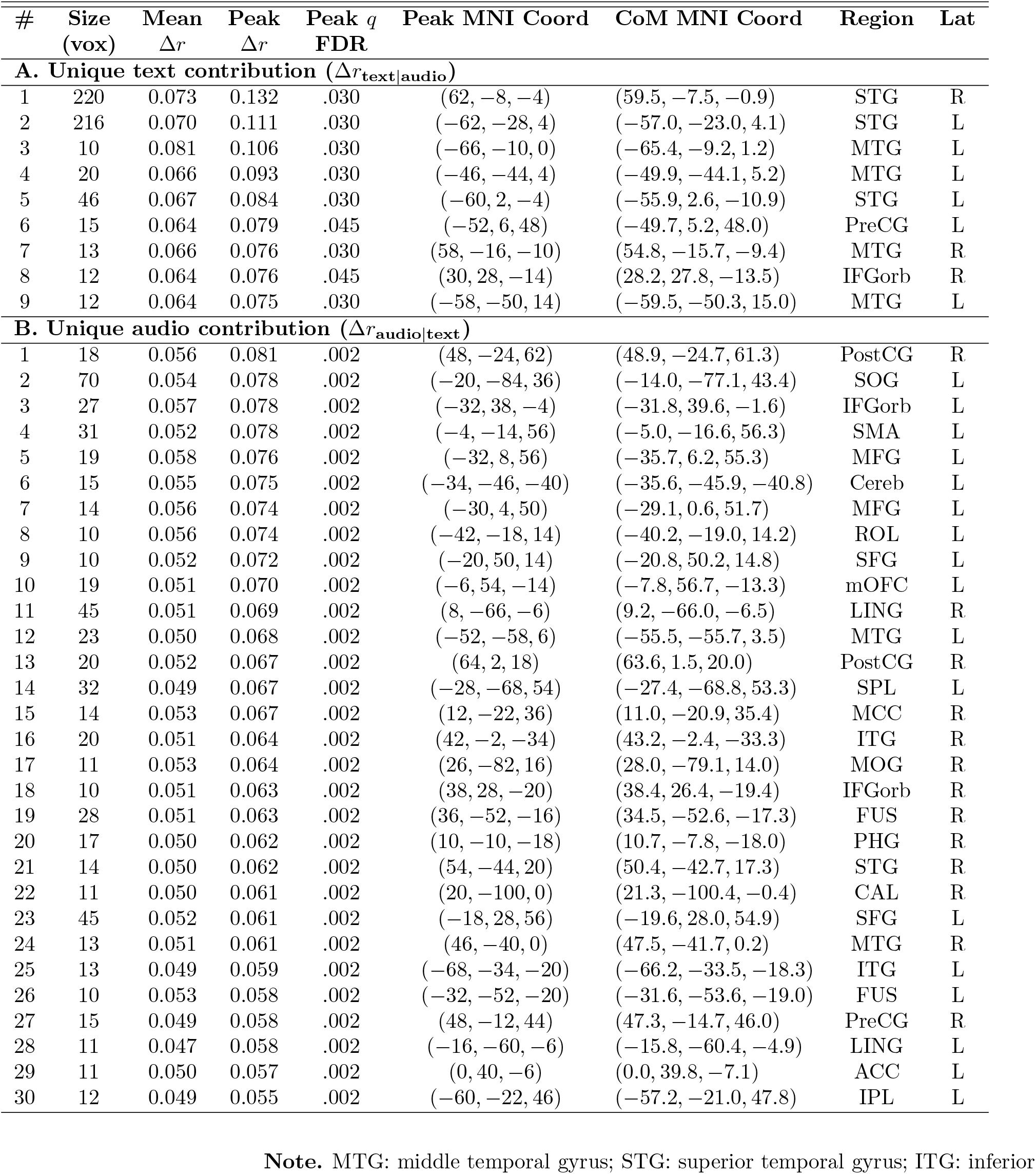
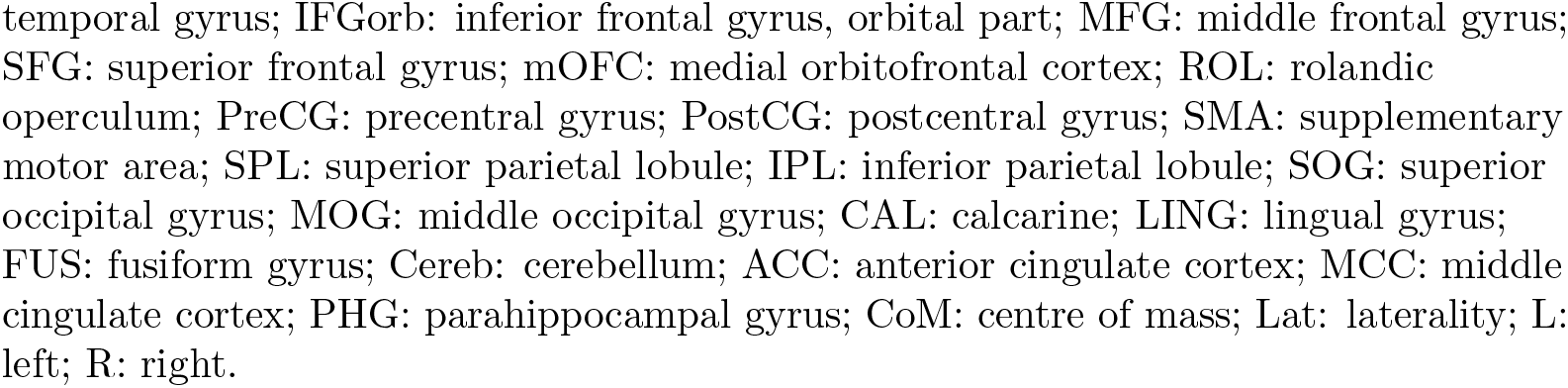
Conditional-contribution clusters underlying the integration conjunction (context-attention embeddings). For the context-attention model, each panel reports the clusters in which one modality contributed unique predictive variance over the other (top 1% of positive Δ*r*, FDR *q <* .05, clusters ≥ 10 contiguous voxels). Panel **A**: unique text contribution Δ*r*_text|audio_; Panel **B**: unique audio contribution Δ*r*_audio|text_. The integration map (Table 1A) retains voxels that were FDR-significant in *both* conditional tests and fell within the top 1% of the conjunction statistic Δ*r*_int_; because each panel is thresholded on its own contribution, an integrative region need not form a displayed cluster in both panels.

| # | Size<br>(vox) | Mean<br>$\Delta r$ | Peak<br>$\Delta r$ | Peak $q$<br>FDR | Peak MNI Coord | CoM MNI Coord | Region | Lat |
| --- | --- | --- | --- | --- | --- | --- | --- | --- |
| **A. Unique text contribution ($\Delta r_{\text{text}\vert \text{audio}}$)** |  |  |  |  |  |  |  |  |
| 1 | 220 | 0.073 | 0.132 | .030 | (62, -8, -4) | (59.5, -7.5, -0.9) | STG | R |
| 2 | 216 | 0.070 | 0.111 | .030 | (-62, -28, 4) | (-57.0, -23.0, 4.1) | STG | L |
| 3 | 10 | 0.081 | 0.106 | .030 | (-66, -10, 0) | (-65.4, -9.2, 1.2) | MTG | L |
| 4 | 20 | 0.066 | 0.093 | .030 | (-46, -44, 4) | (-49.9, -44.1, 5.2) | MTG | L |
| 5 | 46 | 0.067 | 0.084 | .030 | (-60, 2, -4) | (-55.9, 2.6, -10.9) | STG | L |
| 6 | 15 | 0.064 | 0.079 | .045 | (-52, 6, 48) | (-49.7, 5.2, 48.0) | PreCG | L |
| 7 | 13 | 0.066 | 0.076 | .030 | (58, -16, -10) | (54.8, -15.7, -9.4) | MTG | R |
| 8 | 12 | 0.064 | 0.076 | .045 | (30, 28, -14) | (28.2, 27.8, -13.5) | IFGorb | R |
| 9 | 12 | 0.064 | 0.075 | .030 | (-58, -50, 14) | (-59.5, -50.3, 15.0) | MTG | L |
| **B. Unique audio contribution ($\Delta r_{\text{audio}\vert \text{text}}$)** |  |  |  |  |  |  |  |  |
| 1 | 18 | 0.056 | 0.081 | .002 | (48, -24, 62) | (48.9, -24.7, 61.3) | PostCG | R |
| 2 | 70 | 0.054 | 0.078 | .002 | (-20, -84, 36) | (-14.0, -77.1, 43.4) | SOG | L |
| 3 | 27 | 0.057 | 0.078 | .002 | (-32, 38, -4) | (-31.8, 39.6, -1.6) | IFGorb | L |
| 4 | 31 | 0.052 | 0.078 | .002 | (-4, -14, 56) | (-5.0, -16.6, 56.3) | SMA | L |
| 5 | 19 | 0.058 | 0.076 | .002 | (-32, 8, 56) | (-35.7, 6.2, 55.3) | MFG | L |
| 6 | 15 | 0.055 | 0.075 | .002 | (-34, -46, -40) | (-35.6, -45.9, -40.8) | Cereb | L |
| 7 | 14 | 0.056 | 0.074 | .002 | (-30, 4, 50) | (-29.1, 0.6, 51.7) | MFG | L |
| 8 | 10 | 0.056 | 0.074 | .002 | (-42, -18, 14) | (-40.2, -19.0, 14.2) | ROL | L |
| 9 | 10 | 0.052 | 0.072 | .002 | (-20, 50, 14) | (-20.8, 50.2, 14.8) | SFG | L |
| 10 | 19 | 0.051 | 0.070 | .002 | (-6, 54, -14) | (-7.8, 56.7, -13.3) | mOFC | L |
| 11 | 45 | 0.051 | 0.069 | .002 | (8, -66, -6) | (9.2, -66.0, -6.5) | LING | R |
| 12 | 23 | 0.050 | 0.068 | .002 | (-52, -58, 6) | (-55.5, -55.7, 3.5) | MTG | L |
| 13 | 20 | 0.052 | 0.067 | .002 | (64, 2, 18) | (63.6, 1.5, 20.0) | PostCG | R |
| 14 | 32 | 0.049 | 0.067 | .002 | (-28, -68, 54) | (-27.4, -68.8, 53.3) | SPL | L |
| 15 | 14 | 0.053 | 0.067 | .002 | (12, -22, 36) | (11.0, -20.9, 35.4) | MCC | R |
| 16 | 20 | 0.051 | 0.064 | .002 | (42, -2, -34) | (43.2, -2.4, -33.3) | ITG | R |
| 17 | 11 | 0.053 | 0.064 | .002 | (26, -82, 16) | (28.0, -79.1, 14.0) | MOG | R |
| 18 | 10 | 0.051 | 0.063 | .002 | (38, 28, -20) | (38.4, 26.4, -19.4) | IFGorb | R |
| 19 | 28 | 0.051 | 0.063 | .002 | (36, -52, -16) | (34.5, -52.6, -17.3) | FUS | R |
| 20 | 17 | 0.050 | 0.062 | .002 | (10, -10, -18) | (10.7, -7.8, -18.0) | PHG | R |
| 21 | 14 | 0.050 | 0.062 | .002 | (54, -44, 20) | (50.4, -42.7, 17.3) | STG | R |
| 22 | 11 | 0.050 | 0.061 | .002 | (20, -100, 0) | (21.3, -100.4, -0.4) | CAL | R |
| 23 | 45 | 0.052 | 0.061 | .002 | (-18, 28, 56) | (-19.6, 28.0, 54.9) | SFG | L |
| 24 | 13 | 0.051 | 0.061 | .002 | (46, -40, 0) | (47.5, -41.7, 0.2) | MTG | R |
| 25 | 13 | 0.049 | 0.059 | .002 | (-68, -34, -20) | (-66.2, -33.5, -18.3) | ITG | L |
| 26 | 10 | 0.053 | 0.058 | .002 | (-32, -52, -20) | (-31.6, -53.6, -19.0) | FUS | L |
| 27 | 15 | 0.049 | 0.058 | .002 | (48, -12, 44) | (47.3, -14.7, 46.0) | PreCG | R |
| 28 | 11 | 0.047 | 0.058 | .002 | (-16, -60, -6) | (-15.8, -60.4, -4.9) | LING | L |
| 29 | 11 | 0.050 | 0.057 | .002 | (0, 40, -6) | (0.0, 39.8, -7.1) | ACC | L |
| 30 | 12 | 0.049 | 0.055 | .002 | (-60, -22, 46) | (-57.2, -21.0, 47.8) | IPL | L |

**Note.** MTG: middle temporal gyrus; STG: superior temporal gyrus; ITG: inferior
temporal gyrus; IFGorb: inferior frontal gyrus, orbital part; MFG: middle frontal gyrus; SFG: superior frontal gyrus; mOFC: medial orbitofrontal cortex; ROL: rolandic operculum; PreCG: precentral gyrus; PostCG: postcentral gyrus; SMA: supplementary motor area; SPL: superior parietal lobule; IPL: inferior parietal lobule; SOG: superior occipital gyrus; MOG: middle occipital gyrus; CAL: calcarine; LING: lingual gyrus; FUS: fusiform gyrus; Cereb: cerebellum; ACC: anterior cingulate cortex; MCC: middle cingulate cortex; PHG: parahippocampal gyrus; CoM: centre of mass; Lat: laterality; L: left; R: right.

**S2 Fig.**
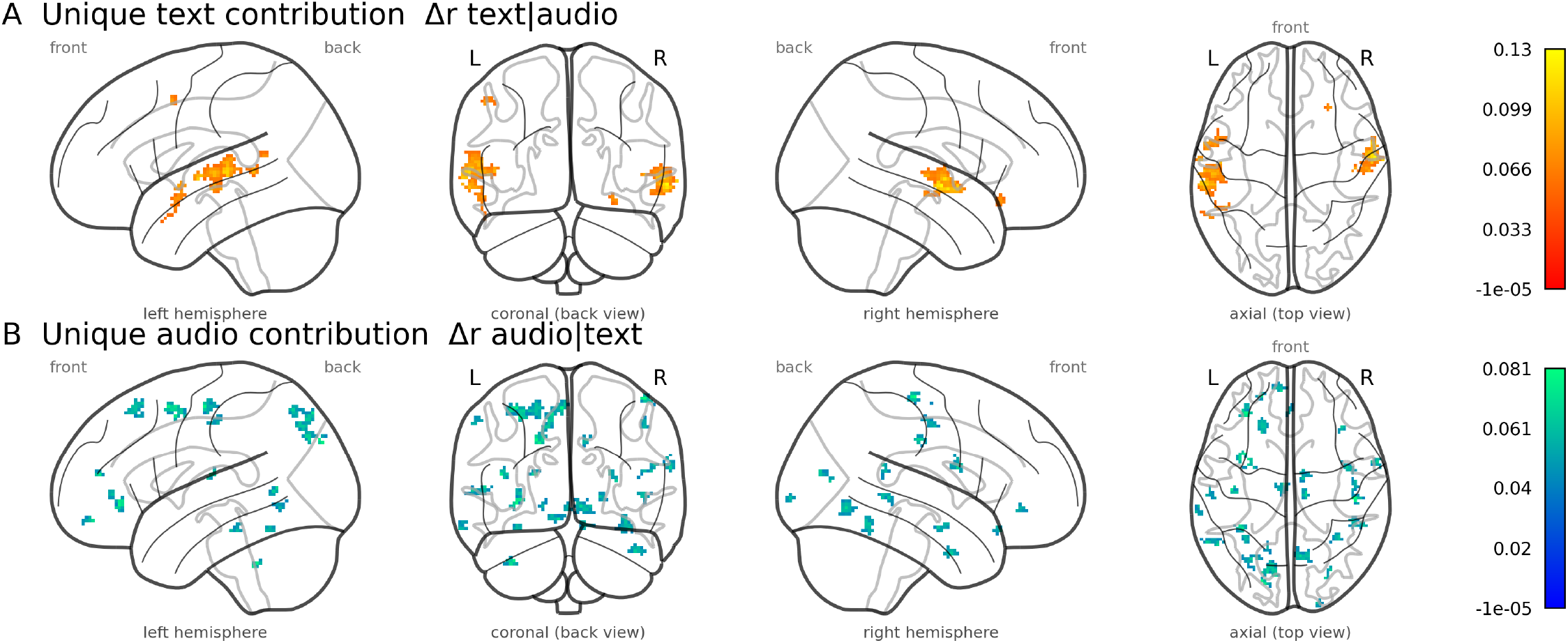
Conditional-contribution maps for the context-attention model. Glass-brain maps of the two conditional contributions whose conjunction defines the integration map (Fig 2A). Panel **A**: unique text contribution Δ*r*_text|audio_ = *r*_text+audio_ − *r*_audio_. Panel **B**: unique audio contribution Δ*r*_audio|text_ = *r*_text+audio_ − *r*_text_. Both maps display voxels surviving the top 1% positive threshold (FDR-corrected *q <* .05, clusters ≥ 10 voxels); cluster details are listed in S1 Table.

**S3 Fig.**
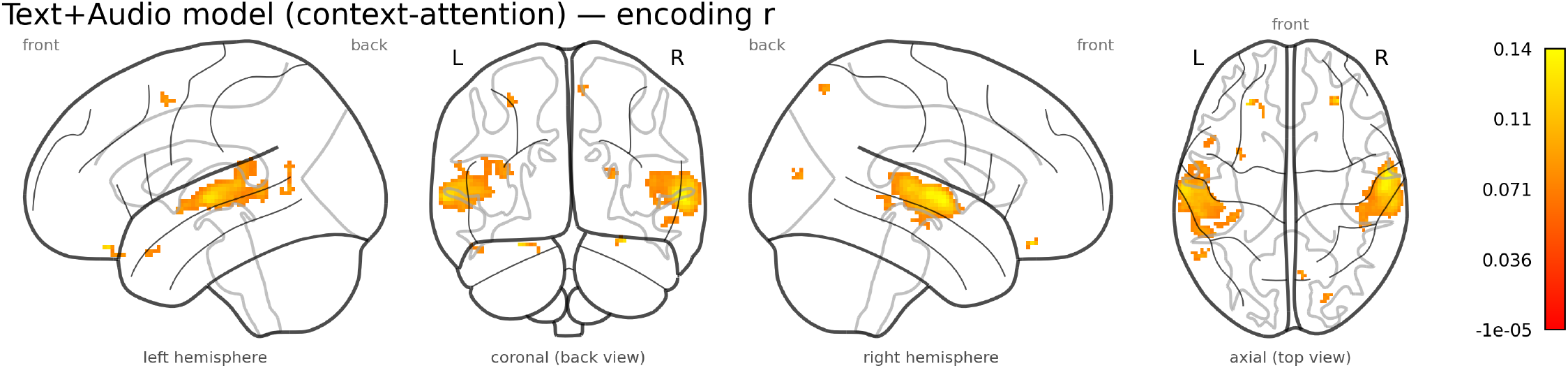
Combined text+audio encoding performance (context-attention embeddings). Glass-brain map of voxel-wise brain scores (Pearson *r*) for the combined text+audio model, surviving the top 1% positive threshold (FDR-corrected *q <* .05, clusters ≥ 10 voxels). The map reproduces the bilateral temporal pattern of the unimodal models (Fig 1).

**S4 Fig.**
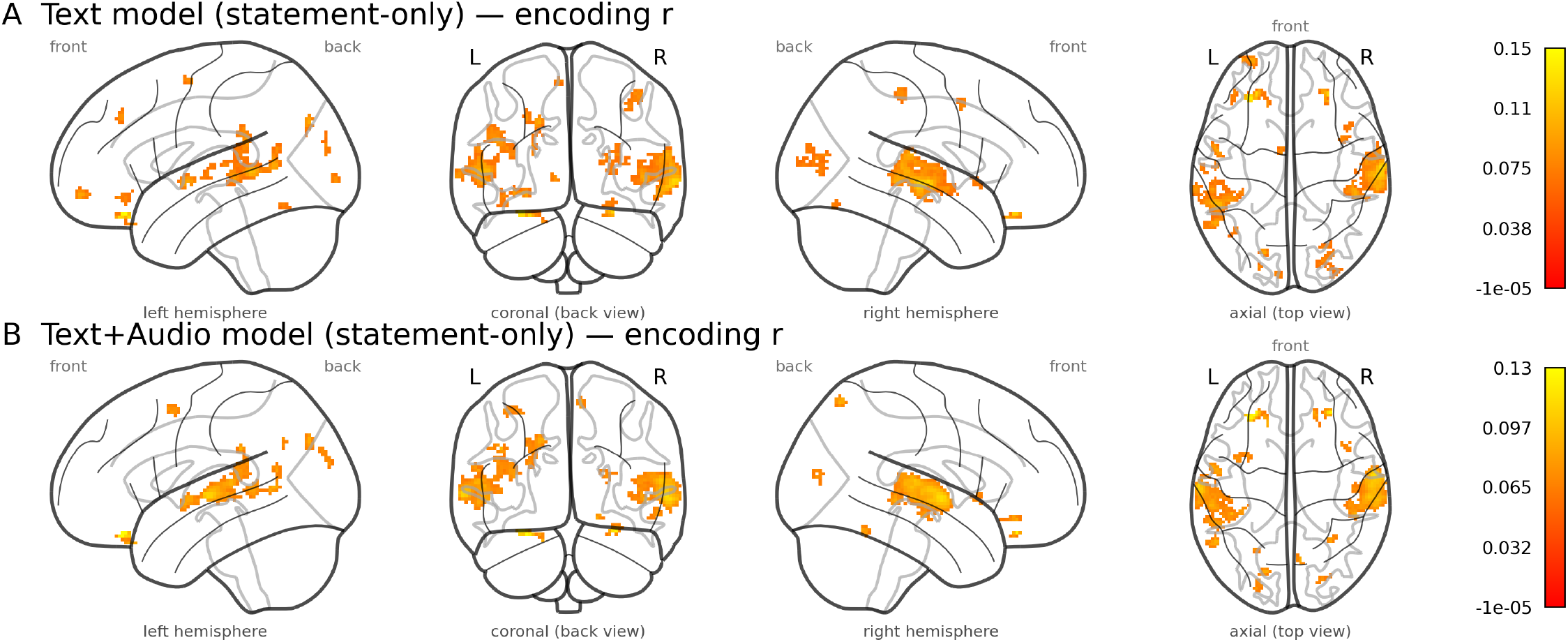
Encoding performance for the statement-only embeddings. Glass-brain maps of voxel-wise brain scores (Pearson *r*) surviving the top 1% positive threshold (FDR-corrected *q <* .05, clusters ≥ 10 voxels). Panel **A**: text model (statement-only embeddings). Panel **B**: combined text+audio model (statement-only embeddings). Cluster details are listed in S6 Table and S7 Table, respectively. These maps are the statement-only counterparts of Fig 1 and S3 Fig.

**S5 Fig.**
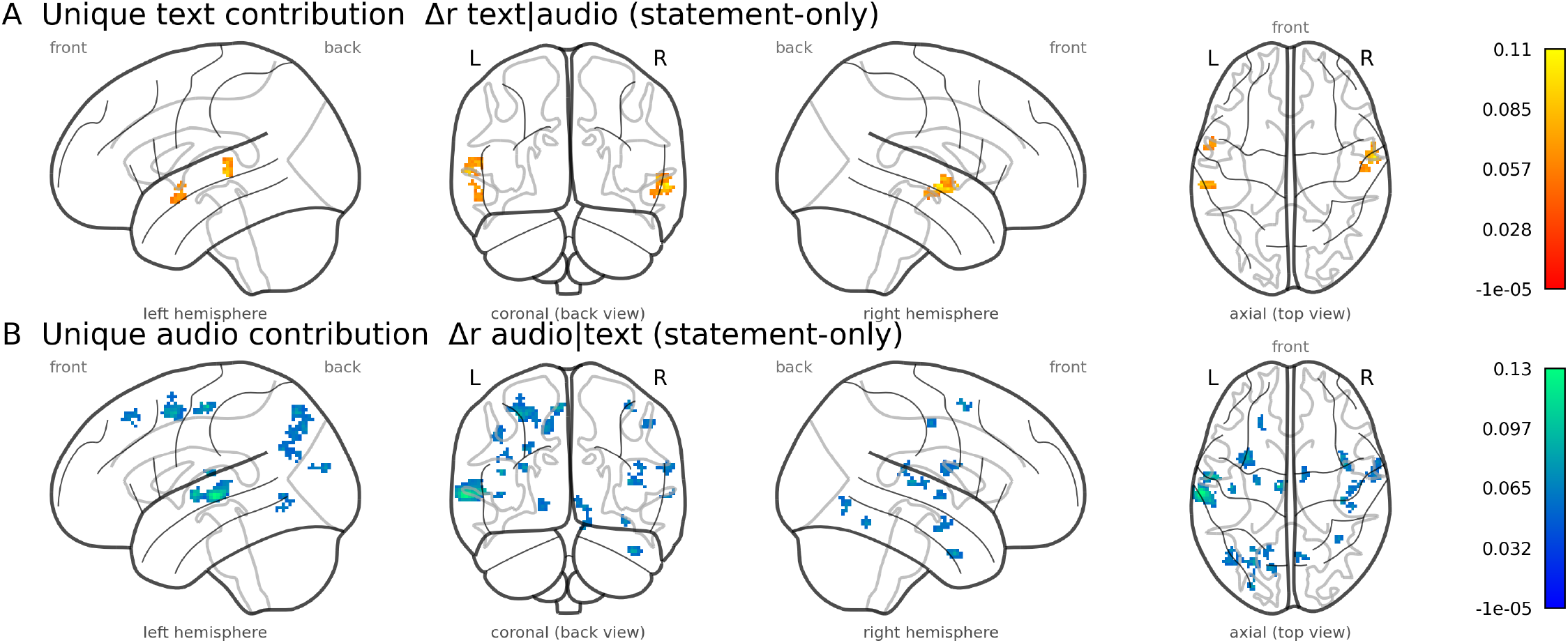
Conditional-contribution maps for the statement-only model. Glass-brain maps of the two conditional contributions whose conjunction defines the statement-only integration map (Fig 2B). Panel **A**: unique text contribution Δ*r*_text|audio_ = *r*_text+audio_ − *r*_audio_. Panel **B**: unique audio contribution Δ*r*_audio|text_ = *r*_text+audio_ − *r*_text_. Both maps display voxels surviving the top 1% positive threshold (FDR-corrected *q <* .05, clusters ≥ 10 voxels); cluster details are listed in S3 Table.

**S2 Table.**
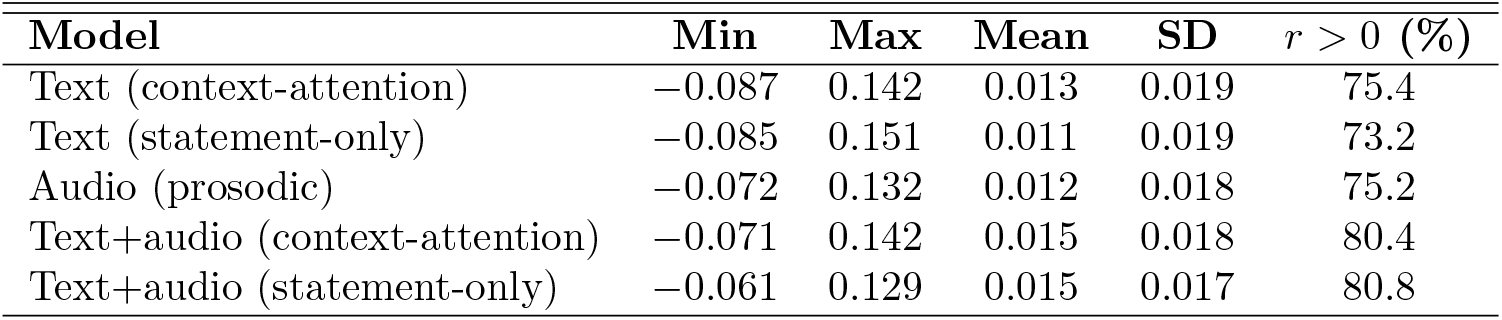
Whole-brain encoding performance of all models. For each model, the distribution of voxel-wise brain scores (Pearson *r* between predicted and observed responses) across in-brain voxels: minimum, maximum, mean, standard deviation, and percentage of voxels with positive scores.

**S3 Table.**
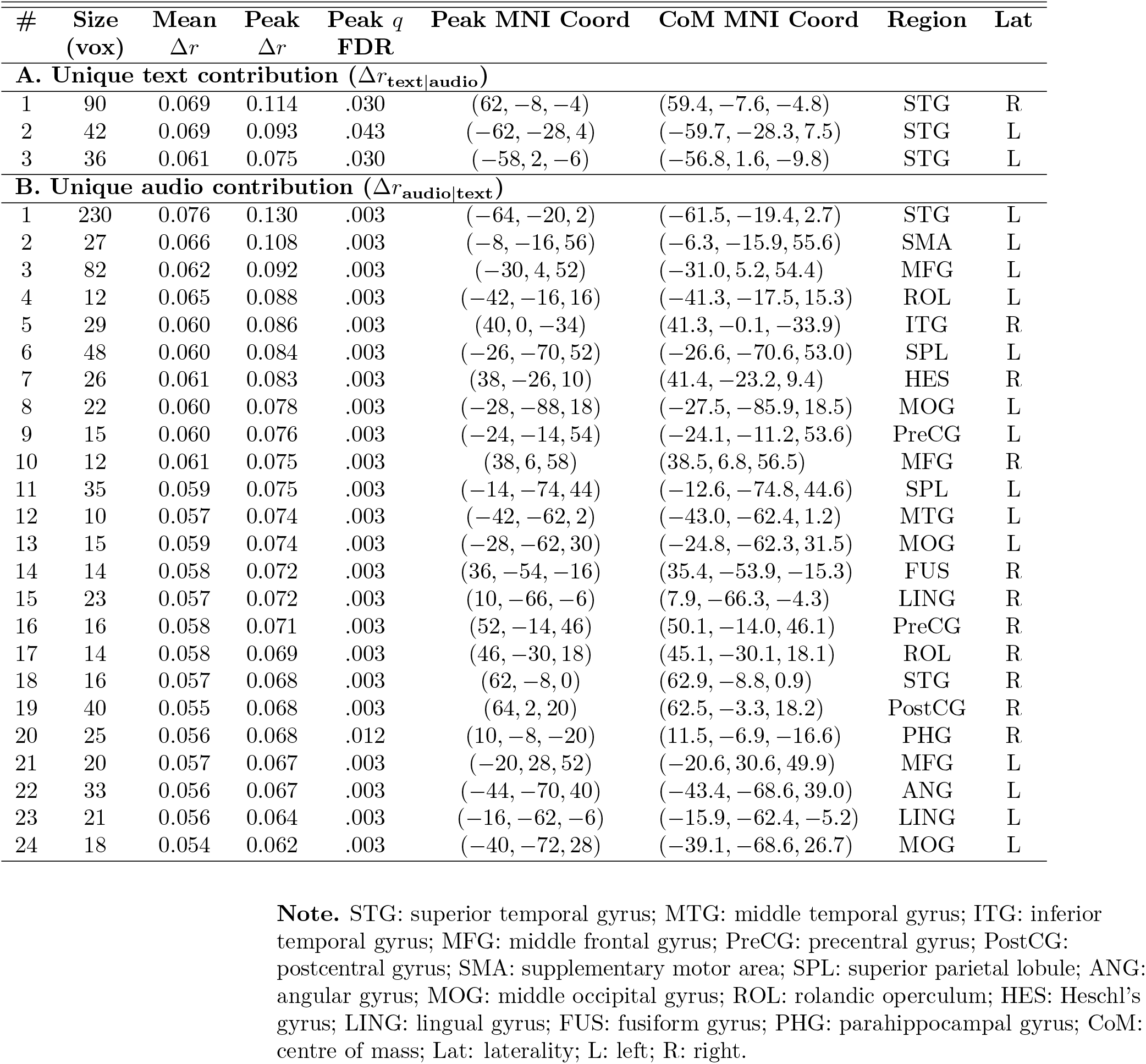
Conditional-contribution clusters underlying the statement-only integration conjunction. For the statement-only model, each panel reports the clusters in which one modality contributed unique predictive variance over the other (top 1% of positive Δ*r*, FDR *q <* .05, clusters ≥ 10 contiguous voxels). Panel **A**: unique text contribution Δ*r*_text|audio_; Panel **B**: unique audio contribution Δ*r*_audio|text_. The statement-only integration map (Table 1B) retains voxels that were FDR-significant in *both* conditional tests and fell within the top 1% of the conjunction statistic. Within each panel, clusters are ranked in descending order of peak Δ*r*.

**S4 Table.** Whole-brain encoding clusters for the text model (context-attention embeddings). Clusters of voxels whose brain score (Pearson *r* between predicted and observed responses) survived the top 1% positive threshold (FDR *q <* .05, clusters ≥ 10 contiguous voxels), ranked in descending order of peak *r*. For each cluster we report its size (in voxels), mean and peak *r*, FDR-corrected *q*-value at the peak voxel, peak and centre-of-mass (CoM) MNI coordinates, anatomical region (AAL atlas), and hemispheric laterality (Lat).

| # | Size<br>(vox) | Mean<br>$r$ | Peak<br>$r$ | Peak $q$<br>FDR | Peak MNI Coord | CoM MNI Coord | Region | Lat |
| --- | --- | --- | --- | --- | --- | --- | --- | --- |
| 1 | 854 | 0.095 | 0.142 | .031 | (60, -8, 0) | (61, -18, 2) | STG | R |
| 2 | 740 | 0.090 | 0.127 | .031 | (-54, -42, 4) | (-57, -25, 3) | MTG | L |
| 3 | 12 | 0.094 | 0.124 | .031 | (28, 38, -22) | (26, 37, -23) | OFCant | R |
| 4 | 10 | 0.083 | 0.098 | .031 | (24, -84, 12) | (23, -83, 12) | SOG | R |
| 5 | 12 | 0.084 | 0.094 | .031 | (-28, 62, -10) | (-28, 61, -11) | SFG | L |
| 6 | 62 | 0.081 | 0.092 | .031 | (-34, -34, 14) | (-41, -37, 22) | RolOp | L |
| 7 | 13 | 0.081 | 0.088 | .031 | (46, -30, 48) | (44, -32, 50) | PostCG | R |

**Note.** STG: superior temporal gyrus; MTG: middle temporal gyrus; SOG: superior occipital gyrus; SFG: superior frontal gyrus; OFCant: anterior orbital gyrus; PostCG: postcentral gyrus; RolOp: rolandic operculum; CoM: centre of mass; Lat: laterality; L: left; R: right.

**S5 Table.** Whole-brain encoding clusters for the audio (prosodic) model. Clusters of voxels whose brain score (Pearson *r*) survived the top 1% positive threshold (FDR *q <* .05, clusters ≥ 10 contiguous voxels), ranked in descending order of peak *r*. Columns as in S4 Table.

| # | Size<br>(vox) | Mean<br>$r$ | Peak<br>$r$ | Peak $q$<br>FDR | Peak MNI Coord | CoM MNI Coord | Region | Lat |
| --- | --- | --- | --- | --- | --- | --- | --- | --- |
| 1 | 42 | 0.083 | 0.132 | .041 | (-30, 34, -22) | (-25, 34, -24) | OFCpost | L |
| 2 | 44 | 0.074 | 0.102 | .041 | (8, -70, 58) | (8, -69, 59) | PCUN | R |
| 3 | 239 | 0.069 | 0.096 | .041 | (-16, -78, 36) | (-20, -83, 31) | CUN | L |
| 4 | 503 | 0.071 | 0.095 | .041 | (58, -24, 10) | (62, -22, 3) | STG | R |
| 5 | 108 | 0.067 | 0.095 | .041 | (-66, -14, 2) | (-63, -18, 1) | STG | L |
| 6 | 23 | 0.068 | 0.094 | .041 | (-44, -36, 30) | (-43, -36, 31) | SMG | L |
| 7 | 32 | 0.070 | 0.090 | .041 | (48, -36, -2) | (48, -36, -2) | MTG | R |
| 8 | 25 | 0.069 | 0.087 | .041 | (-24, -62, 32) | (-25, -63, 33) | MOG | L |
| 9 | 20 | 0.069 | 0.085 | .041 | (-54, -56, 6) | (-54, -55, 5) | MTG | L |
| 10 | 15 | 0.068 | 0.084 | .041 | (-32, 8, 54) | (-34, 7, 55) | MFG | L |
| 11 | 30 | 0.068 | 0.083 | .041 | (28, -56, -20) | (30, -54, -21) | Cereb | R |
| 12 | 48 | 0.066 | 0.082 | .041 | (-36, 60, -14) | (-33, 60, -12) | MFG | L |
| 13 | 31 | 0.068 | 0.082 | .041 | (-52, -4, -2) | (-52, -5, -1) | STG | L |
| 14 | 53 | 0.067 | 0.081 | .041 | (30, -78, 10) | (25, -82, 13) | MOG | R |
| 15 | 26 | 0.066 | 0.079 | .041 | (-56, -58, 16) | (-57, -58, 17) | MTG | L |
| 16 | 35 | 0.067 | 0.078 | .041 | (38, 4, 44) | (40, 5, 48) | MFG | R |
| 17 | 34 | 0.064 | 0.077 | .041 | (-58, -56, 30) | (-57, -56, 28) | SMG | L |
| 18 | 109 | 0.065 | 0.077 | .041 | (18, -78, 28) | (22, -76, 33) | SOG | R |
| 19 | 12 | 0.066 | 0.075 | .041 | (-62, -10, -18) | (-62, -8, -19) | MTG | L |
| 20 | 21 | 0.064 | 0.071 | .041 | (-38, -64, -22) | (-39, -64, -18) | Cereb | L |
| 21 | 22 | 0.064 | 0.070 | .041 | (14, 20, 56) | (14, 19, 56) | SMA | R |
| 22 | 12 | 0.062 | 0.068 | .041 | (-38, -70, 28) | (-39, -68, 27) | MOG | L |

**Note.** STG: superior temporal gyrus; MTG: middle temporal gyrus; MFG: middle frontal gyrus; OFCpost: posterior orbital gyrus; SMA: supplementary motor area; SMG: supramarginal gyrus; MOG: middle occipital gyrus; SOG: superior occipital gyrus; CUN: cuneus; PCUN: precuneus; Cereb: cerebellum; CoM: centre of mass; Lat: laterality; L: left; R: right.

**S6 Table.** Whole-brain encoding clusters for the text model (statement-only embeddings). Clusters of voxels whose brain score (Pearson *r*) survived the top 1% positive threshold (FDR *q <* .05, clusters ≥ 10 contiguous voxels), ranked in descending order of peak *r*. Columns as in S4 Table.

| # | Size<br>(vox) | Mean<br>$r$ | Peak<br>$r$ | Peak $q$<br>FDR | Peak MNI Coord | CoM MNI Coord | Region | Lat |
| --- | --- | --- | --- | --- | --- | --- | --- | --- |
| 1 | 43 | 0.093 | 0.151 | .034 | (−30, 34, −22) | (−23, 35, −24) | OFCpost | L |
| 2 | 579 | 0.082 | 0.128 | .034 | (66, −16, −2) | (62, −21, 2) | STG | R |
| 3 | 19 | 0.089 | 0.124 | .034 | (28, 36, −22) | (26, 36, −23) | OFCant | R |
| 4 | 318 | 0.078 | 0.112 | .034 | (−54, −42, 4) | (−56, −43, 5) | MTG | L |
| 5 | 95 | 0.078 | 0.102 | .034 | (48, −34, −2) | (48, −34, 2) | MTG | R |
| 6 | 28 | 0.078 | 0.101 | .034 | (−18, −78, 34) | (−18, −77, 33) | CUN | L |
| 7 | 36 | 0.080 | 0.099 | .034 | (−52, −4, −2) | (−54, −3, −2) | STG | L |
| 8 | 48 | 0.077 | 0.098 | .034 | (−28, 62, −10) | (−29, 61, −10) | SFG | L |
| 9 | 103 | 0.075 | 0.096 | .034 | (−44, −40, 24) | (−42, −38, 22) | SMG | L |
| 10 | 18 | 0.074 | 0.087 | .034 | (−26, 36, 38) | (−25, 37, 37) | SFG | L |
| 11 | 15 | 0.075 | 0.087 | .034 | (−8, −94, −2) | (−8, −94, −1) | CAL | L |
| 12 | 13 | 0.074 | 0.085 | .034 | (−6, −6, 60) | (−6, −4, 59) | SMA | L |
| 13 | 61 | 0.071 | 0.084 | .034 | (24, −84, 12) | (27, −84, 11) | SOG | R |
| 14 | 14 | 0.072 | 0.084 | .034 | (−22, −88, 20) | (−21, −88, 21) | MOG | L |
| 15 | 36 | 0.074 | 0.084 | .034 | (42, −34, 52) | (43, −32, 51) | PostCG | R |
| 16 | 23 | 0.071 | 0.083 | .034 | (−40, 38, −12) | (−41, 35, −12) | IFGorb | L |
| 17 | 23 | 0.069 | 0.081 | .034 | (−56, −26, 8) | (−50, −29, 10) | STG | L |
| 18 | 20 | 0.072 | 0.080 | .034 | (38, 4, 44) | (39, 4, 46) | MFG | R |
| 19 | 11 | 0.072 | 0.079 | .034 | (−52, −16, 2) | (−54, −15, 1) | STG | L |
| 20 | 14 | 0.071 | 0.078 | .034 | (−38, −62, −16) | (−40, −63, −17) | FUS | L |
| 21 | 15 | 0.071 | 0.077 | .034 | (26, −74, −18) | (27, −73, −17) | Cereb | R |
| 22 | 10 | 0.071 | 0.077 | .034 | (−64, −16, 4) | (−62, −17, 6) | STG | L |
| 23 | 15 | 0.069 | 0.076 | .034 | (34, −92, 8) | (34, −93, 9) | MOG | R |
| 24 | 17 | 0.069 | 0.074 | .034 | (44, 14, −2) | (43, 13, −2) | INS | R |

**Note.** STG: superior temporal gyrus; MTG: middle temporal gyrus; SFG: superior frontal gyrus; MFG: middle frontal gyrus; OFCpost: posterior orbital gyrus; OFCant: anterior orbital gyrus; IFGorb: inferior frontal gyrus, orbital part; PostCG: postcentral gyrus; SMA: supplementary motor area; SMG: supramarginal gyrus; MOG: middle occipital gyrus; SOG: superior occipital gyrus; CUN: cuneus; CAL: calcarine; FUS: fusiform gyrus; Cereb: cerebellum; INS: insula; CoM: centre of mass; Lat: laterality; L: left; R: right.

**S7 Table.** Whole-brain encoding clusters for the combined text+audio model (statement-only embeddings). Clusters of voxels whose brain score (Pearson *r*) survived the top 1% positive threshold (FDR *q <* .05, clusters ≥ 10 contiguous voxels), ranked in descending order of peak *r*. Columns as in S4 Table.

| # | Size<br>(vox) | Mean<br>$r$ | Peak<br>$r$ | Peak $q$<br>FDR | Peak MNI Coord | CoM MNI Coord | Region | Lat |
| --- | --- | --- | --- | --- | --- | --- | --- | --- |
| 1 | 37 | 0.089 | 0.129 | .029 | (−26, 36, −24) | (−23, 35, −24) | OFCant | L |
| 2 | 11 | 0.084 | 0.123 | .029 | (30, 38, −22) | (28, 38, −22) | OFCant | R |
| 3 | 830 | 0.076 | 0.113 | .029 | (62, −8, 0) | (60, −20, 3) | STG | R |
| 4 | 450 | 0.072 | 0.108 | .029 | (−66, −16, 0) | (−55, −26, 8) | MTG | L |
| 5 | 17 | 0.073 | 0.091 | .029 | (8, −70, 58) | (8, −70, 59) | PCUN | R |
| 6 | 123 | 0.070 | 0.089 | .029 | (−54, −56, 6) | (−54, −48, 7) | MTG | L |
| 7 | 36 | 0.070 | 0.088 | .029 | (−16, −78, 36) | (−18, −78, 34) | CUN | L |
| 8 | 15 | 0.070 | 0.086 | .029 | (18, 38, −14) | (19, 35, −13) | OFCmed | R |
| 9 | 51 | 0.073 | 0.085 | .029 | (−56, −2, −4) | (−54, −4, −3) | STG | L |
| 10 | 18 | 0.070 | 0.082 | .029 | (30, −56, −20) | (31, −54, −20) | Cereb | R |
| 11 | 23 | 0.068 | 0.079 | .029 | (56, −18, −10) | (55, −18, −12) | MTG | R |
| 12 | 30 | 0.067 | 0.077 | .029 | (−24, −90, 20) | (−23, −88, 24) | MOG | L |
| 13 | 32 | 0.067 | 0.076 | .029 | (−30, 6, 52) | (−35, 5, 53) | MFG | L |
| 14 | 11 | 0.067 | 0.075 | .029 | (−26, −62, 36) | (−24, −62, 34) | MOG | L |
| 15 | 19 | 0.065 | 0.074 | .029 | (24, −84, 12) | (24, −83, 13) | SOG | R |
| 16 | 16 | 0.066 | 0.072 | .029 | (38, 14, 2) | (41, 13, 0) | INS | R |

**Note.** STG: superior temporal gyrus; MTG: middle temporal gyrus; MFG: middle frontal gyrus; OFCant: anterior orbital gyrus; OFCmed: medial orbital gyrus; MOG: middle occipital gyrus; SOG: superior occipital gyrus; CUN: cuneus; PCUN: precuneus; INS: insula; Cereb: cerebellum; CoM: centre of mass; Lat: laterality; L: left; R: right.

**S8 Table.** Whole-brain encoding clusters for the combined text+audio model (context-attention embeddings). Clusters of voxels whose brain score (Pearson *r*) survived the top 1% positive threshold (FDR *q <* .05, clusters ≥ 10 contiguous voxels), ranked in descending order of peak *r*. Columns as in S4 Table. This is the combined-model counterpart of the map shown in S3 Fig.

| # | Size<br>(vox) | Mean<br>$r$ | Peak<br>$r$ | Peak $q$<br>FDR | Peak MNI Coord | CoM MNI Coord | Region | Lat |
| --- | --- | --- | --- | --- | --- | --- | --- | --- |
| 1 | 922 | 0.091 | 0.142 | .030 | (62, −8, 0) | (60, −19, 3) | STG | R |
| 2 | 750 | 0.085 | 0.127 | .030 | (−64, −18, 2) | (−57, −23, 4) | STG | L |
| 3 | 10 | 0.091 | 0.126 | .030 | (30, 38, −22) | (28, 38, −22) | OFCant | R |
| 4 | 13 | 0.088 | 0.125 | .030 | (−26, 36, −24) | (−20, 35, −25) | OFCant | L |
| 5 | 23 | 0.078 | 0.094 | .030 | (−54, −58, 6) | (−55, −58, 10) | MTG | L |
| 6 | 10 | 0.078 | 0.091 | .030 | (8, −70, 58) | (8, −70, 59) | PCUN | R |
| 7 | 13 | 0.078 | 0.090 | .030 | (−30, 6, 52) | (−31, 6, 52) | MFG | L |
| 8 | 54 | 0.076 | 0.090 | .030 | (−34, −34, 14) | (−37, −35, 17) | RolOp | L |
| 9 | 11 | 0.077 | 0.089 | .030 | (24, −82, 12) | (23, −84, 14) | CAL | R |
| 10 | 16 | 0.074 | 0.082 | .030 | (52, −24, −10) | (55, −19, −11) | MTG | R |
| 11 | 13 | 0.074 | 0.081 | .030 | (−48, 16, −28) | (−48, 13, −26) | TP | L |

**Note.** STG: superior temporal gyrus; MTG: middle temporal gyrus; MFG: middle frontal gyrus; OFCant: anterior orbital gyrus; PCUN: precuneus; CAL: calcarine; RolOp: rolandic operculum; TP: temporal pole; CoM: centre of mass; Lat: laterality; L: left; R: right.

**S9 Table.**
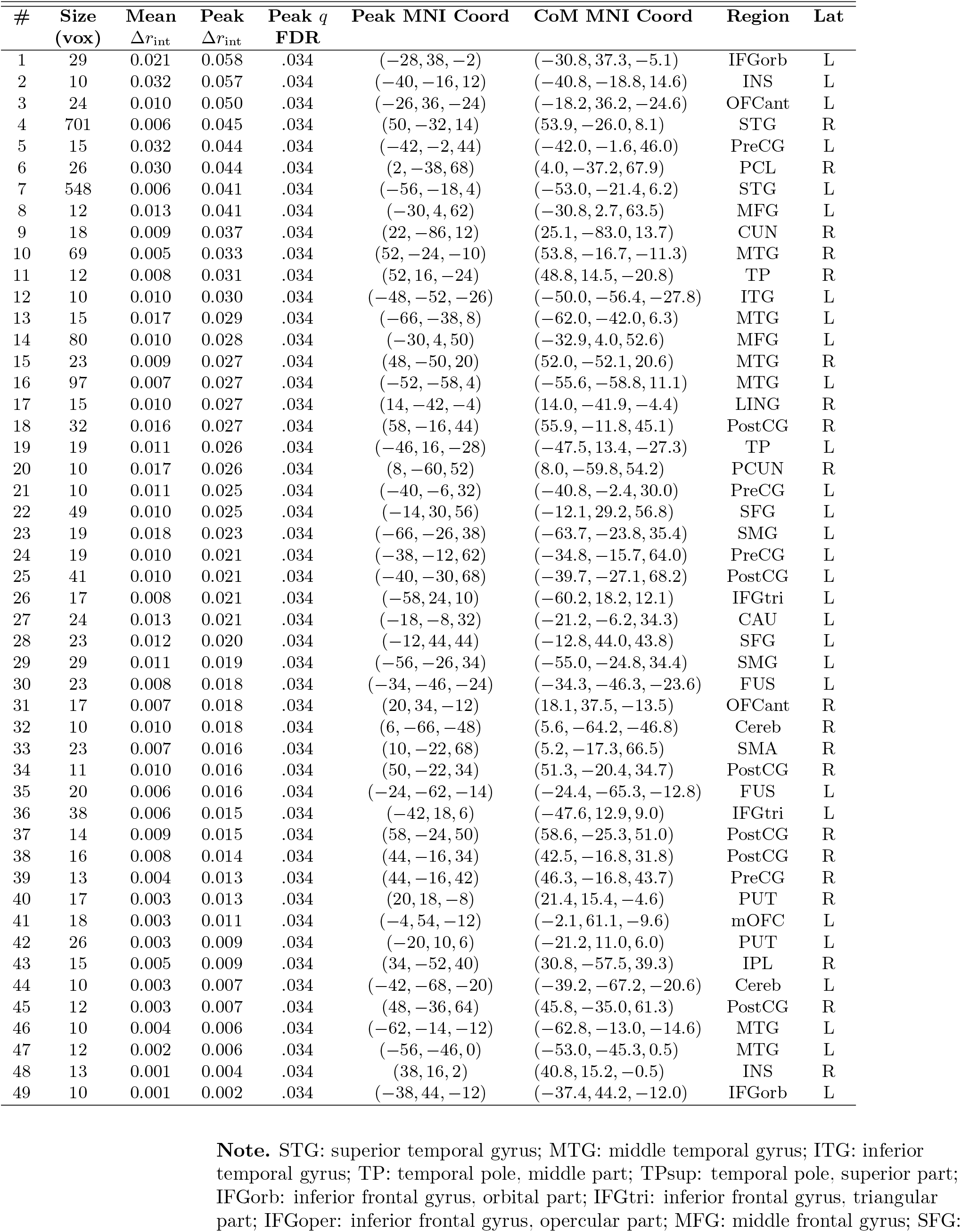
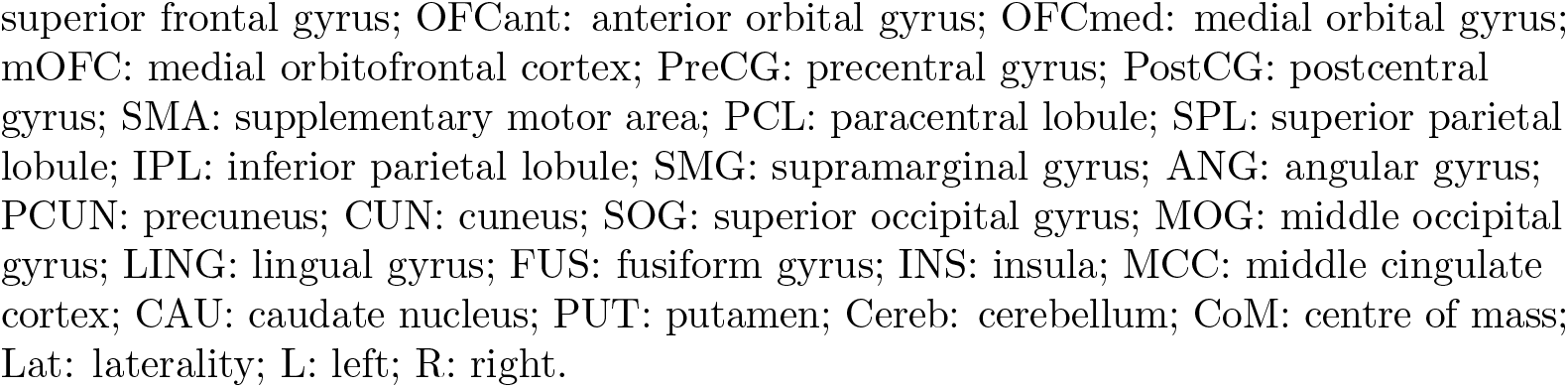
Integration clusters without the percentile threshold (context-attention embeddings). Clusters in which *both* conditional contributions (Δ*r*_audio|text_ and Δ*r*_text|audio_) were positive and FDR-significant (*q <* .05) and that formed clusters of at least 10 contiguous voxels, *without* the additional top 1% restriction applied in Table 1. Columns are as in Table 1; clusters are ranked in descending order of peak Δ*r*_int_. Peak *q*-values are identical across clusters because the permutation *p*-values are discrete (*p* = (*b* + 1)*/*(*m* + 1), *m* = 1,000) and every cluster peak attains the smallest possible value. Orbital regions are labelled using AAL3, as in S4 Table–S8 Table.

| # | Size<br>(vox) | Mean<br>$\Delta r_{\text{int}}$ | Peak<br>$\Delta r_{\text{int}}$ | Peak $q$<br>FDR | Peak MNI Coord | CoM MNI Coord | Region | Lat |
| --- | --- | --- | --- | --- | --- | --- | --- | --- |
| 1 | 29 | 0.021 | 0.058 | .034 | (-28, 38, -2) | (-30.8, 37.3, -5.1) | IFGorb | L |
| 2 | 10 | 0.032 | 0.057 | .034 | (-40, -16, 12) | (-40.8, -18.8, 14.6) | INS | L |
| 3 | 24 | 0.010 | 0.050 | .034 | (-26, 36, -24) | (-18.2, 36.2, -24.6) | OFCant | L |
| 4 | 701 | 0.006 | 0.045 | .034 | (50, -32, 14) | (53.9, -26.0, 8.1) | STG | R |
| 5 | 15 | 0.032 | 0.044 | .034 | (-42, -2, 44) | (-42.0, -1.6, 46.0) | PreCG | L |
| 6 | 26 | 0.030 | 0.044 | .034 | (2, -38, 68) | (4.0, -37.2, 67.9) | PCL | R |
| 7 | 548 | 0.006 | 0.041 | .034 | (-56, -18, 4) | (-53.0, -21.4, 6.2) | STG | L |
| 8 | 12 | 0.013 | 0.041 | .034 | (-30, 4, 62) | (-30.8, 2.7, 63.5) | MFG | L |
| 9 | 18 | 0.009 | 0.037 | .034 | (22, -86, 12) | (25.1, -83.0, 13.7) | CUN | R |
| 10 | 69 | 0.005 | 0.033 | .034 | (52, -24, -10) | (53.8, -16.7, -11.3) | MTG | R |
| 11 | 12 | 0.008 | 0.031 | .034 | (52, 16, -24) | (48.8, 14.5, -20.8) | TP | R |
| 12 | 10 | 0.010 | 0.030 | .034 | (-48, -52, -26) | (-50.0, -56.4, -27.8) | ITG | L |
| 13 | 15 | 0.017 | 0.029 | .034 | (-66, -38, 8) | (-62.0, -42.0, 6.3) | MTG | L |
| 14 | 80 | 0.010 | 0.028 | .034 | (-30, 4, 50) | (-32.9, 4.0, 52.6) | MFG | L |
| 15 | 23 | 0.009 | 0.027 | .034 | (48, -50, 20) | (52.0, -52.1, 20.6) | MTG | R |
| 16 | 97 | 0.007 | 0.027 | .034 | (-52, -58, 4) | (-55.6, -58.8, 11.1) | MTG | L |
| 17 | 15 | 0.010 | 0.027 | .034 | (14, -42, -4) | (14.0, -41.9, -4.4) | LING | R |
| 18 | 32 | 0.016 | 0.027 | .034 | (58, -16, 44) | (55.9, -11.8, 45.1) | PostCG | R |
| 19 | 19 | 0.011 | 0.026 | .034 | (-46, 16, -28) | (-47.5, 13.4, -27.3) | TP | L |
| 20 | 10 | 0.017 | 0.026 | .034 | (8, -60, 52) | (8.0, -59.8, 54.2) | PCUN | R |
| 21 | 10 | 0.011 | 0.025 | .034 | (-40, -6, 32) | (-40.8, -2.4, 30.0) | PreCG | L |
| 22 | 49 | 0.010 | 0.025 | .034 | (-14, 30, 56) | (-12.1, 29.2, 56.8) | SFG | L |
| 23 | 19 | 0.018 | 0.023 | .034 | (-66, -26, 38) | (-63.7, -23.8, 35.4) | SMG | L |
| 24 | 19 | 0.010 | 0.021 | .034 | (-38, -12, 62) | (-34.8, -15.7, 64.0) | PreCG | L |
| 25 | 41 | 0.010 | 0.021 | .034 | (-40, -30, 68) | (-39.7, -27.1, 68.2) | PostCG | L |
| 26 | 17 | 0.008 | 0.021 | .034 | (-58, 24, 10) | (-60.2, 18.2, 12.1) | IFGtri | L |
| 27 | 24 | 0.013 | 0.021 | .034 | (-18, -8, 32) | (-21.2, -6.2, 34.3) | CAU | L |
| 28 | 23 | 0.012 | 0.020 | .034 | (-12, 44, 44) | (-12.8, 44.0, 43.8) | SFG | L |
| 29 | 29 | 0.011 | 0.019 | .034 | (-56, -26, 34) | (-55.0, -24.8, 34.4) | SMG | L |
| 30 | 23 | 0.008 | 0.018 | .034 | (-34, -46, -24) | (-34.3, -46.3, -23.6) | FUS | L |
| 31 | 17 | 0.007 | 0.018 | .034 | (20, 34, -12) | (18.1, 37.5, -13.5) | OFCant | R |
| 32 | 10 | 0.010 | 0.018 | .034 | (6, -66, -48) | (5.6, -64.2, -46.8) | Cereb | R |
| 33 | 23 | 0.007 | 0.016 | .034 | (10, -22, 68) | (5.2, -17.3, 66.5) | SMA | R |
| 34 | 11 | 0.010 | 0.016 | .034 | (50, -22, 34) | (51.3, -20.4, 34.7) | PostCG | R |
| 35 | 20 | 0.006 | 0.016 | .034 | (-24, -62, -14) | (-24.4, -65.3, -12.8) | FUS | L |
| 36 | 38 | 0.006 | 0.015 | .034 | (-42, 18, 6) | (-47.6, 12.9, 9.0) | IFGtri | L |
| 37 | 14 | 0.009 | 0.015 | .034 | (58, -24, 50) | (58.6, -25.3, 51.0) | PostCG | R |
| 38 | 16 | 0.008 | 0.014 | .034 | (44, -16, 34) | (42.5, -16.8, 31.8) | PostCG | R |
| 39 | 13 | 0.004 | 0.013 | .034 | (44, -16, 42) | (46.3, -16.8, 43.7) | PreCG | R |
| 40 | 17 | 0.003 | 0.013 | .034 | (20, 18, -8) | (21.4, 15.4, -4.6) | PUT | R |
| 41 | 18 | 0.003 | 0.011 | .034 | (-4, 54, -12) | (-2.1, 61.1, -9.6) | mOFC | L |
| 42 | 26 | 0.003 | 0.009 | .034 | (-20, 10, 6) | (-21.2, 11.0, 6.0) | PUT | L |
| 43 | 15 | 0.005 | 0.009 | .034 | (34, -52, 40) | (30.8, -57.5, 39.3) | IPL | R |
| 44 | 10 | 0.003 | 0.007 | .034 | (-42, -68, -20) | (-39.2, -67.2, -20.6) | Cereb | L |
| 45 | 12 | 0.003 | 0.007 | .034 | (48, -36, 64) | (45.8, -35.0, 61.3) | PostCG | R |
| 46 | 10 | 0.004 | 0.006 | .034 | (-62, -14, -12) | (-62.8, -13.0, -14.6) | MTG | L |
| 47 | 12 | 0.002 | 0.006 | .034 | (-56, -46, 0) | (-53.0, -45.3, 0.5) | MTG | L |
| 48 | 13 | 0.001 | 0.004 | .034 | (38, 16, 2) | (40.8, 15.2, -0.5) | INS | R |
| 49 | 10 | 0.001 | 0.002 | .034 | (-38, 44, -12) | (-37.4, 44.2, -12.0) | IFGorb | L |

**Note.** STG: superior temporal gyrus; MTG: middle temporal gyrus; ITG: inferior temporal gyrus; TP: temporal pole, middle part; TPsup: temporal pole, superior part; IFGorb: inferior frontal gyrus, orbital part; IFGtri: inferior frontal gyrus, triangular part; IFGoper: inferior frontal gyrus, opercular part; MFG: middle frontal gyrus; SFG:
superior frontal gyrus; OFCant: anterior orbital gyrus; OFCmed: medial orbital gyrus; 909 mOFC: medial orbitofrontal cortex; PreCG: precentral gyrus; PostCG: postcentral 910 gyrus; SMA: supplementary motor area; PCL: paracentral lobule; SPL: superior parietal 911 lobule; IPL: inferior parietal lobule; SMG: supramarginal gyrus; ANG: angular gyrus; 912 PCUN: precuneus; CUN: cuneus; SOG: superior occipital gyrus; MOG: middle occipital 913 gyrus; LING: lingual gyrus; FUS: fusiform gyrus; INS: insula; MCC: middle cingulate 914 cortex; CAU: caudate nucleus; PUT: putamen; Cereb: cerebellum; CoM: centre of mass; 915 Lat: laterality; L: left; R: right. 916

**S10 Table.**
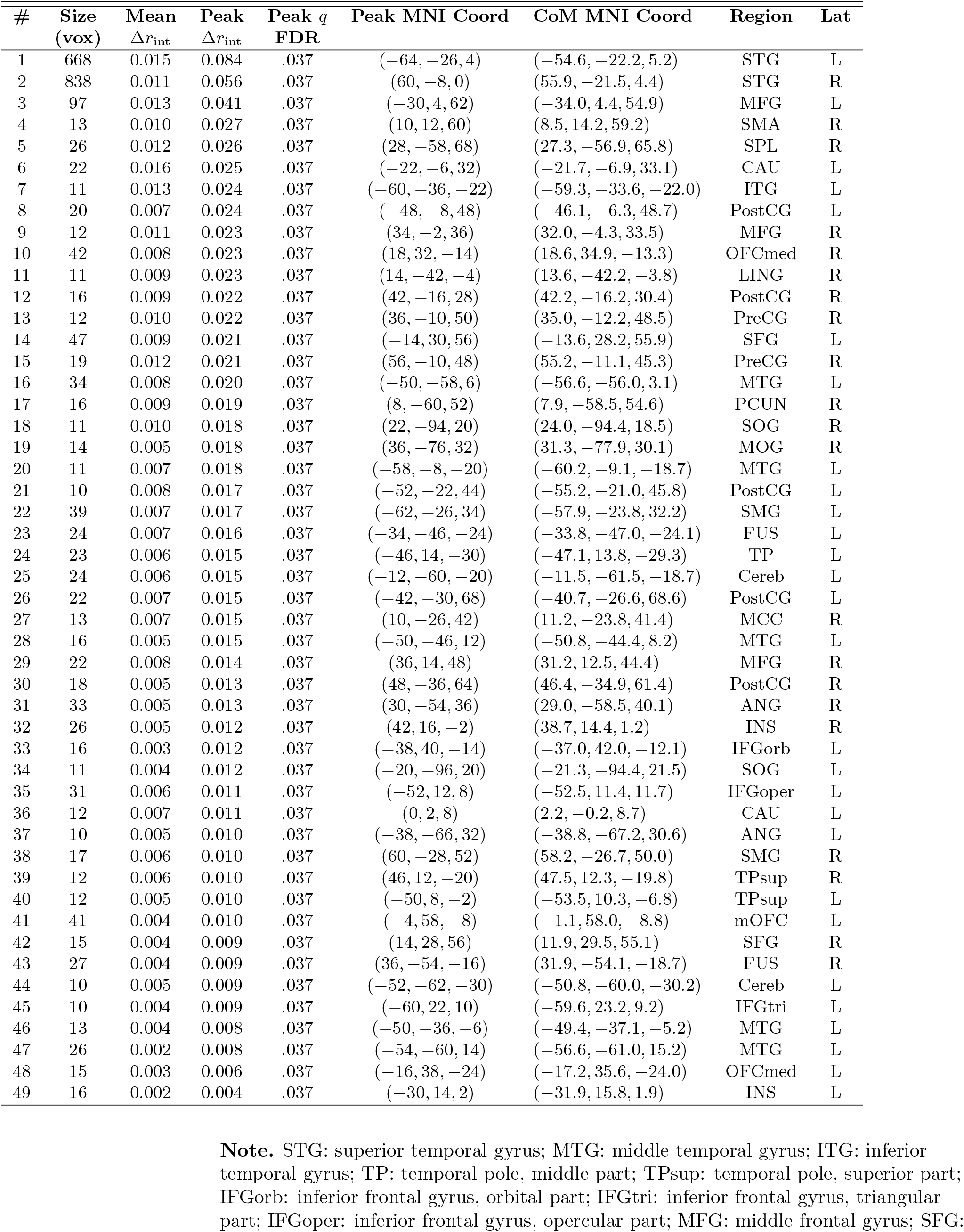
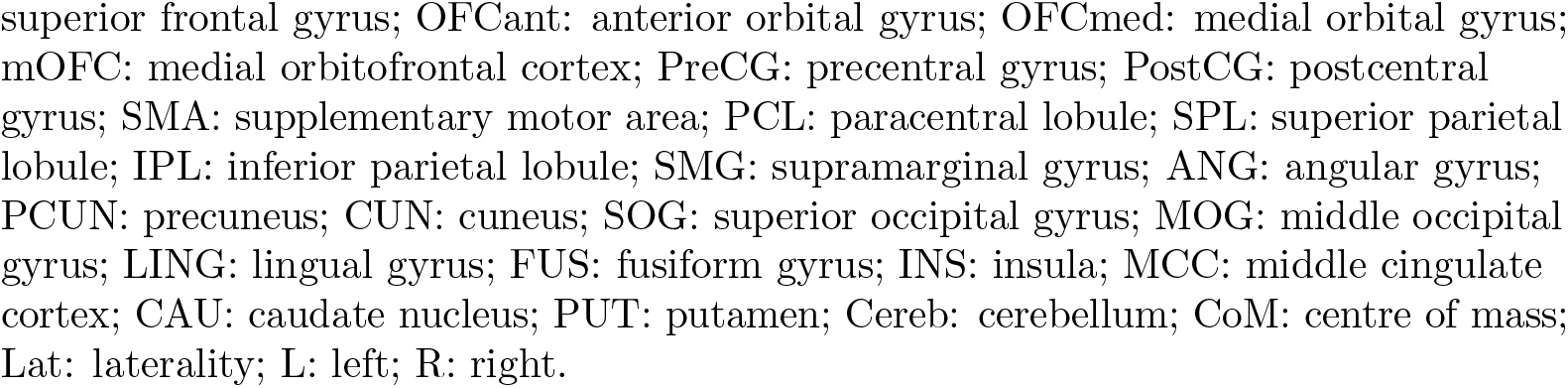
Integration clusters without the percentile threshold (statement-only embeddings). Clusters in which *both* conditional contributions (Δ*r*_audio|text_ and Δ*r*_text|audio_) were positive and FDR-significant (*q <* .05) and that formed clusters of at least 10 contiguous voxels, *without* the additional top 1% restriction applied in Table 1. Columns are as in Table 1; clusters are ranked in descending order of peak Δ*r*_int_. Peak *q*-values are identical across clusters because the permutation *p*-values are discrete (*p* = (*b* + 1)*/*(*m* + 1), *m* = 1,000) and every cluster peak attains the smallest possible value. Orbital regions are labelled using AAL3, as in S4 Table–S8 Table.

| # | Size<br>(vox) | Mean<br>$\Delta r_{\text{int}}$ | Peak<br>$\Delta r_{\text{int}}$ | Peak $q$<br>FDR | Peak MNI Coord | CoM MNI Coord | Region | Lat |
| --- | --- | --- | --- | --- | --- | --- | --- | --- |
| 1 | 668 | 0.015 | 0.084 | .037 | (-64, -26, 4) | (-54.6, -22.2, 5.2) | STG | L |
| 2 | 838 | 0.011 | 0.056 | .037 | (60, -8, 0) | (55.9, -21.5, 4.4) | STG | R |
| 3 | 97 | 0.013 | 0.041 | .037 | (-30, 4, 62) | (-34.0, 4.4, 54.9) | MFG | L |
| 4 | 13 | 0.010 | 0.027 | .037 | (10, 12, 60) | (8.5, 14.2, 59.2) | SMA | R |
| 5 | 26 | 0.012 | 0.026 | .037 | (28, -58, 68) | (27.3, -56.9, 65.8) | SPL | R |
| 6 | 22 | 0.016 | 0.025 | .037 | (-22, -6, 32) | (-21.7, -6.9, 33.1) | CAU | L |
| 7 | 11 | 0.013 | 0.024 | .037 | (-60, -36, -22) | (-59.3, -33.6, -22.0) | ITG | L |
| 8 | 20 | 0.007 | 0.024 | .037 | (-48, -8, 48) | (-46.1, -6.3, 48.7) | PostCG | L |
| 9 | 12 | 0.011 | 0.023 | .037 | (34, -2, 36) | (32.0, -4.3, 33.5) | MFG | R |
| 10 | 42 | 0.008 | 0.023 | .037 | (18, 32, -14) | (18.6, 34.9, -13.3) | OFCmed | R |
| 11 | 11 | 0.009 | 0.023 | .037 | (14, -42, -4) | (13.6, -42.2, -3.8) | LING | R |
| 12 | 16 | 0.009 | 0.022 | .037 | (42, -16, 28) | (42.2, -16.2, 30.4) | PostCG | R |
| 13 | 12 | 0.010 | 0.022 | .037 | (36, -10, 50) | (35.0, -12.2, 48.5) | PreCG | R |
| 14 | 47 | 0.009 | 0.021 | .037 | (-14, 30, 56) | (-13.6, 28.2, 55.9) | SFG | L |
| 15 | 19 | 0.012 | 0.021 | .037 | (56, -10, 48) | (55.2, -11.1, 45.3) | PreCG | R |
| 16 | 34 | 0.008 | 0.020 | .037 | (-50, -58, 6) | (-56.6, -56.0, 3.1) | MTG | L |
| 17 | 16 | 0.009 | 0.019 | .037 | (8, -60, 52) | (7.9, -58.5, 54.6) | PCUN | R |
| 18 | 11 | 0.010 | 0.018 | .037 | (22, -94, 20) | (24.0, -94.4, 18.5) | SOG | R |
| 19 | 14 | 0.005 | 0.018 | .037 | (36, -76, 32) | (31.3, -77.9, 30.1) | MOG | R |
| 20 | 11 | 0.007 | 0.018 | .037 | (-58, -8, -20) | (-60.2, -9.1, -18.7) | MTG | L |
| 21 | 10 | 0.008 | 0.017 | .037 | (-52, -22, 44) | (-55.2, -21.0, 45.8) | PostCG | L |
| 22 | 39 | 0.007 | 0.017 | .037 | (-62, -26, 34) | (-57.9, -23.8, 32.2) | SMG | L |
| 23 | 24 | 0.007 | 0.016 | .037 | (-34, -46, -24) | (-33.8, -47.0, -24.1) | FUS | L |
| 24 | 23 | 0.006 | 0.015 | .037 | (-46, 14, -30) | (-47.1, 13.8, -29.3) | TP | L |
| 25 | 24 | 0.006 | 0.015 | .037 | (-12, -60, -20) | (-11.5, -61.5, -18.7) | Cereb | L |
| 26 | 22 | 0.007 | 0.015 | .037 | (-42, -30, 68) | (-40.7, -26.6, 68.6) | PostCG | L |
| 27 | 13 | 0.007 | 0.015 | .037 | (10, -26, 42) | (11.2, -23.8, 41.4) | MCC | R |
| 28 | 16 | 0.005 | 0.015 | .037 | (-50, -46, 12) | (-50.8, -44.4, 8.2) | MTG | L |
| 29 | 22 | 0.008 | 0.014 | .037 | (36, 14, 48) | (31.2, 12.5, 44.4) | MFG | R |
| 30 | 18 | 0.005 | 0.013 | .037 | (48, -36, 64) | (46.4, -34.9, 61.4) | PostCG | R |
| 31 | 33 | 0.005 | 0.013 | .037 | (30, -54, 36) | (29.0, -58.5, 40.1) | ANG | R |
| 32 | 26 | 0.005 | 0.012 | .037 | (42, 16, -2) | (38.7, 14.4, 1.2) | INS | R |
| 33 | 16 | 0.003 | 0.012 | .037 | (-38, 40, -14) | (-37.0, 42.0, -12.1) | IFGorb | L |
| 34 | 11 | 0.004 | 0.012 | .037 | (-20, -96, 20) | (-21.3, -94.4, 21.5) | SOG | L |
| 35 | 31 | 0.006 | 0.011 | .037 | (-52, 12, 8) | (-52.5, 11.4, 11.7) | IFGoper | L |
| 36 | 12 | 0.007 | 0.011 | .037 | (0, 2, 8) | (2.2, -0.2, 8.7) | CAU | L |
| 37 | 10 | 0.005 | 0.010 | .037 | (-38, -66, 32) | (-38.8, -67.2, 30.6) | ANG | L |
| 38 | 17 | 0.006 | 0.010 | .037 | (60, -28, 52) | (58.2, -26.7, 50.0) | SMG | R |
| 39 | 12 | 0.006 | 0.010 | .037 | (46, 12, -20) | (47.5, 12.3, -19.8) | TPsup | R |
| 40 | 12 | 0.005 | 0.010 | .037 | (-50, 8, -2) | (-53.5, 10.3, -6.8) | TPsup | L |
| 41 | 41 | 0.004 | 0.010 | .037 | (-4, 58, -8) | (-1.1, 58.0, -8.8) | mOFC | L |
| 42 | 15 | 0.004 | 0.009 | .037 | (14, 28, 56) | (11.9, 29.5, 55.1) | SFG | R |
| 43 | 27 | 0.004 | 0.009 | .037 | (36, -54, -16) | (31.9, -54.1, -18.7) | FUS | R |
| 44 | 10 | 0.005 | 0.009 | .037 | (-52, -62, -30) | (-50.8, -60.0, -30.2) | Cereb | L |
| 45 | 10 | 0.004 | 0.009 | .037 | (-60, 22, 10) | (-59.6, 23.2, 9.2) | IFGtri | L |
| 46 | 13 | 0.004 | 0.008 | .037 | (-50, -36, -6) | (-49.4, -37.1, -5.2) | MTG | L |
| 47 | 26 | 0.002 | 0.008 | .037 | (-54, -60, 14) | (-56.6, -61.0, 15.2) | MTG | L |
| 48 | 15 | 0.003 | 0.006 | .037 | (-16, 38, -24) | (-17.2, 35.6, -24.0) | OFCmed | L |
| 49 | 16 | 0.002 | 0.004 | .037 | (-30, 14, 2) | (-31.9, 15.8, 1.9) | INS | L |

**Note.** STG: superior temporal gyrus; MTG: middle temporal gyrus; ITG: inferior temporal gyrus; TP: temporal pole, middle part; TPsup: temporal pole, superior part; IFGorb: inferior frontal gyrus, orbital part; IFGtri: inferior frontal gyrus, triangular part; IFGoper: inferior frontal gyrus, opercular part; MFG: middle frontal gyrus; SFG:
superior frontal gyrus; OFCant: anterior orbital gyrus; OFCmed: medial orbital gyrus; 930 mOFC: medial orbitofrontal cortex; PreCG: precentral gyrus; PostCG: postcentral 931 gyrus; SMA: supplementary motor area; PCL: paracentral lobule; SPL: superior parietal 932 lobule; IPL: inferior parietal lobule; SMG: supramarginal gyrus; ANG: angular gyrus; 933 PCUN: precuneus; CUN: cuneus; SOG: superior occipital gyrus; MOG: middle occipital 934 gyrus; LING: lingual gyrus; FUS: fusiform gyrus; INS: insula; MCC: middle cingulate 935 cortex; CAU: caudate nucleus; PUT: putamen; Cereb: cerebellum; CoM: centre of mass; 936 Lat: laterality; L: left; R: right. 937

